# Trans-omic analysis reveals fed and fasting insulin signal across phosphoproteome, transcriptome, and metabolome

**DOI:** 10.1101/187088

**Authors:** Kentaro Kawata, Katsuyuki Yugi, Atsushi Hatano, Masashi Fujii, Yoko Tomizawa, Toshiya Kokaji, Takanori Sano, Kaori Y. Tanaka, Shinsuke Uda, Hiroyuki Kubota, Yutaka Suzuki, Masaki Matsumoto, Keiichi I. Nakayama, Kaori Saitoh, Keiko Kato, Ayano Ueno, Maki Ohishi, Tomoyoshi Soga, Shinya Kuroda

**Affiliations:** Department of Biological Sciences, Graduate School of Science, University of Tokyo, 7-3-1 Hongo, Bunkyo-ku, Tokyo 113-0033, Japan; YCI Laboratory for Trans-Omics, Young Chief Investigator Program, RIKEN Center for Integrative Medical Science, 1-7-22 Suehiro-cho, Tsurumi-ku, Yokohama, Kanagawa 230-0045, Japan; Institute for Advanced Biosciences, Keio University, Fujisawa, 252-8520, Japan; Molecular Genetics Research Laboratory, Graduate School of Science, University of Tokyo, 7-3-1 Hongo, Bunkyo-ku, Tokyo 113-0033, Japan; Department of Computational Biology and Medical Sciences, Graduate School of Frontier Sciences, University of Tokyo, 5-1-5 Kashiwanoha, Kashiwa, Chiba 277-8562, Japan; Division of Integrated Omics, Research Center for Transomics Medicine, Medical Institute of Bioregulation, Kyushu University, 3-1-1 Maidashi, Higashi-ku, Fukuoka 812-8582, Japan; Department of Molecular and Cellular Biology, Medical Institute of Bioregulation, Kyushu University, 3-1-1 Maidashi, Higashi-ku, Fukuoka 812-8582, Japan; Institute for Advanced Biosciences, Keio University, 246-2 Mizukami, Kakuganji, Tsuruoka, Yamagata 997-0052, Japan; PRESTO, Japan Science and Technology Agency, 1-7-22 Suehiro-cho, Tsurumi-ku, Yokohama, Kanagawa 230-0045, Japan; Core Research for Evolutional Science and Technology (CREST), Japan Science and Technology Agency, Bunkyo-ku, Tokyo 113-0033, Japan

**Keywords:** Trans-omics, phosphoproteome, transcriptome, metabolome, insulin signaling, gene expression, metabolism, systems biology, network integration

## Abstract

The concentration and temporal pattern of insulin selectively regulate multiple cellular functions. To understand how insulin dynamics are interpreted by cells, we constructed a trans-omic network of insulin action in FAO hepatoma cells from three networks—a phosphorylation-dependent cellular functions regulatory network using phosphoproteomic data, a transcriptional regulatory network using phosphoproteomic and transcriptomic data, and a metabolism regulatory network using phosphoproteomic and metabolomic data. With the trans-omic regulatory network, we identified selective regulatory networks that mediate differential responses to insulin. Akt and Erk, hub molecules of insulin signaling, encode information of a wide dynamic range of dose and time of insulin. Down-regulated genes and metabolites in glycolysis had high sensitivity to insulin (fasting insulin signal); up-regulated genes and dicarboxylic acids in the TCA cycle had low sensitivity (fed insulin signal). This integrated analysis enables molecular insight into how cells interpret physiologically fed and fasting insulin signals.

**Highlights:**

- We constructed a trans-omic network of insulin action using multi-omic data.
- The trans-omic network integrates phosphorylation, transcription, and metabolism.
- We classified signaling, transcriptome, and metabolome by sensitivity to insulin.
- We identified fed and fasting insulin signal flow across the trans-omic network.

## INTRODUCTION

Metabolic disorders involving insulin resistance are a major health concern (Zimmet et al., 2001). Understanding how cells interpret this physiologically dynamic hormone may provide new insights into preventing or treating metabolic disorders associated with insulin resistance. As with many hormones, not only does the release of the hormone vary but the cellular response is complex and changes over time (Brabant et al., 1992; Lindsay et al., 2003; O’Meara et al., 1993; O’Rahilly et al., 1988; Polonsky et al., 1988). Insulin affects enzyme and protein activity, cellular metabolite composition, and gene expression (Lizcano and Alessi, 2002; Saltiel and Kahn, 2001). Individually, each of these can be studied with existing technologies, but a challenge is integrating disparate types of omic data to generate a more comprehensive view of the cellular response than can be gained from one type of data alone (Yugi and Kuroda, 2017; Yugi et al., 2016). Here, we describe a process for integrating networks constructed from phosphoproteomic, transcriptomic, and metabolomic data to produce a trans-omic regulatory network to explore how cells interpret a physiologically dynamic stimulus, insulin.

Insulin regulates organismal metabolic homeostasis by regulating multiple cellular functions, including gene expression, metabolism, and protein synthesis, in target organs such as the liver, skeletal muscle, and adipose tissue (Jastrzebski et al., 2007; Saltiel and Kahn, 2001; Whiteman et al., 2002). Insulin activates the insulin receptor (InsR) (Lizcano and Alessi, 2002; Saltiel and Kahn, 2001). The activated InsR triggers a phosphorylation-mediated signaling pathway that includes the kinases phosphatidylinositol 3-kinase (PI3k), Akt, mTOR complex 1 (mTORC1), extracellular signal-regulated kinases 1 and 2 (Erk1 and Erk2), p38, and adenosine monophosphate-activated protein kinase (Ampk) (Asnaghi et al., 2004; Kotzka et al., 2004; Lizcano and Alessi, 2002; Saltiel and Kahn, 2001; Zhang et al., 2011). Adaptors involved in insulin signaling include insulin receptor substrate 1 and 2 (Irs1 and Irs2) (Lizcano and Alessi, 2002; Saltiel and Kahn, 2001) and growth factor receptor-binding protein 2 (Grb2) (Karoor et al., 1998; Skolnik et al., 1993a, 1993b).

In the liver, insulin-mediated regulation of gene expression mainly occurs through the Akt signaling pathway and the mitogen-activated protein kinase (MAPK) signaling pathways mediated by Erk proteins and p38 (Au et al., 2003; Barthel et al., 2005; Lizcano and Alessi, 2002; Mounier and Posner, 2006; Saltiel and Kahn, 2001). Akt phosphorylates the Foxo family of forkhead transcription factors (TFs) including Foxo1 (also known as Fkhr). Phosphorylated Foxo1 is excluded from the nucleus (Biggs et al., 1999; Brunet et al., 1999), thus Akt-mediated phosphorylation suppresses gluconeogenesis by reducing the expression of *glucose-6-phosphatase* (*G6Pase*) and *phosphoenolpyruvate carboxykinase1* (*Pck1*), which encode rate-limiting enzymes of gluconeogenesis (Lu et al., 2012; Saltiel and Kahn, 2001). Erk regulates the expression of genes, such as *c-Jun*, by activating cAMP response element-binding protein (Creb) and by inducing the protein expression of TFs, such early growth response 1 (Egr1) and hairy and enhancer of split-1 (Hes1) (Deak et al., 1998; Murphy et al., 2004; Nakayama et al., 2008; Shaul and Seger, 2007). c-Jun N-terminal kinase (Jnk) is activated by phosphatidylinositol 3-kinase (PI3k), and Jnk also activates TFs, such as c-Jun and activating transcription factor 2 (Atf2), through phosphorylation (Fukunaga et al., 2000; Lee et al., 2003).

In addition to regulating the expression of genes involved in metabolism, insulin regulates organismal and cellular metabolism, including glycolysis, gluconeogenesis, glycogenesis, amino acid metabolism, and lipid metabolism, by stimulating the phosphorylation of metabolic enzymes. For example, insulin inhibits glycogen synthase kinase 3β (Gsk3β), which inhibits glycogen synthase (GS), thereby promoting glycogen synthesis (Aiston et al., 2006; Srivastava and Pandey, 1998; Whiteman et al., 2002). Insulin also stimulates protein synthesis by activating Akt, which inhibits tuberous sclerosis complex 1 and 2 (Tsc1 and Tsc2), thereby activating mTORC1. Active mTORC1 stimulates protein translation by phosphorylating ribosomal protein S6 kinase (S6k), eukaryotic translation initiation factor 4E-binding protein 1 (eIf4ebp1), and eukaryotic translation initiation factor 4B (eIf4b) (Asnaghi et al., 2004; Humphrey et al., 2013; Rosner et al., 2010).

Insulin regulates other cellular processes, such as cell adhesion, cytoskeletal organization, and RNA splicing (Hartmann et al., 2009; Reiss et al., 2001; Tsakiridis et al., 1999; Wolf et al., 2013). The cellular response to insulin involves changes in posttranslational modifications, allosteric regulation of metabolic enzymes, and changes in gene expression (Whiteman et al., 2002). The coordinated regulation by insulin of all of these cellular functions and processes through these diverse mechanisms involves the transmission of signals across a large-scale network that involves changes in protein phosphorylation, metabolites, mRNAs, and protein synthesis. The trans-omic regulatory networks that enables cells to achieve this coordinated response to insulin has yet to be uncovered.

Technical advances in phosphoproteomic, transcriptomic, and metabolomic measurement enable the large-scale quantitative analysis of changes in protein phosphorylation, transcript abundance, and metabolite abundance. Phosphoproteomic studies of insulin signaling have been performed with *Drosophila* cells (Friedman et al., 2011; Vinayagam et al., 2016), 3T3-L1 mouse adipocytes (Humphrey et al., 2013), mouse hepatoma cells (Monetti et al., 2011), mouse brown preadipocytes (Krüger et al., 2008), rat primary hepatocytes (Zhang et al., 2017), rat FAO hepatoma cells (Yugi et al., 2014), and mouse hepatoma and human cervical cancer cells (Humphrey et al., 2015). Transcriptomic studies of insulin signaling have been performed with rat hepatoma cells (Hectors et al., 2012), mouse fibroblasts (Dupont et al., 2001; Versteyhe et al., 2013), mouse osteoclast precursors (Kim and Lee, 2014), human skeletal muscle (Rome et al., 2003), and FAO cells (Sano et al., 2016). Metabolomic studies of insulin signaling have been performed with human plasma (Everman et al., 2016), and rat FAO hepatoma cells (Noguchi et al., 2013; Yugi et al., 2014). Despite such progress in the individual omic layers of insulin action, little has been reported for interactions between the omic layers (Buescher et al., 2012; Chiappino-Pepe et al., 2017; Hatzimanikatis and Saez-Rodriguez, 2015; Hyduke et al., 2013; Joyce and Palsson, 2006; Palsson and Zengler, 2010).

We propose “trans-omics” as a discipline for constructing molecular interaction networks across multiple omic data sets using direct molecular interactions rather than indirect statistical relationships (Yugi and Kuroda, 2017; Yugi et al., 2014, 2016). Trans-omic analyses of networks controlling metabolism have been reported for *Escherichia coli* (Gerosa et al., 2015; Ishii et al., 2007), *Bacillus subtilis* (Buescher et al., 2012), *Saccharomyces cerevisiae* (Gonçalves et al., 2017; Hackett et al., 2016; Oliveira et al., 2012), Chinese hamster ovary cells (Yusufi et al., 2017), and human T cells (Geiger et al., 2016). We have previously constructed trans-omic networks of the regulation of metabolism through phosphorylation in response to acute insulin action with phosphoproteomic and metabolomic data (Yugi et al., 2014). However, the networks that regulate other cellular functions and the inclusion of data other than phosphoproteomic and metabolomic remain to be constructed.

Temporal patterns of specific growth factors coding extracellular information are encoded into signaling pathways and are selectively decoded by distinct downstream molecules by differences in the sensitivity, time constants, and network structures (Behar and Hoffmann, 2010; Purvis and Lahav, 2013). Glucose stimulates the secretion of insulin from the pancreas resulting in a transient high concentration of insulin in the blood (fed insulin signal); whereas under basal conditions a sustained low concentration of insulin (fasting insulin signal) is maintained in the blood (Lindsay et al., 2003; Polonsky et al., 1988). To respond properly to insulin, cells must be able to detect both fed and fasting insulin signals and interpret each type of insulin signal. We previously showed that signaling molecules such as Akt (Kubota et al., 2012), metabolites such as glycogen (Noguchi et al., 2013), and gene expression such as *G6Pase* and *Pck1* (Sano et al., 2016) show distinct selectivity to a transient high concentration and sustained low concentration of insulin. However, how fed and fasting insulin signals selectively regulate the trans-omic network has yet to be analyzed.

Here, we describe the construction of a trans-omic regulatory network using phosphoproteomic, transcriptomic, and metabolomic data from insulin-stimulated FAO cells. We first constructed three networks: a phosphorylation-dependent cellular functions regulatory network, a transcriptional regulatory network, a metabolism regulatory network involving allosteric regulation and phosphorylation-mediated regulation of metabolic enzyme activity. We integrated these networks to obtain a trans-omic regulatory network that spans the phosphoproteomic, transcriptomic, and metabolomic layers and reveals their regulatory connections. We also estimated sensitivities and time constants of selected signaling molecules, transcriptomic data, and metabolomic data to insulin stimulation, and identified the selective use of specific pathways by transient high concentrations of insulin and low sustained concentration of insulin, reflecting the physiological states of fed and fasting. Thus, our study uses the cellular response to insulin to demonstrate the process of constructing a trans-omic regulatory network and the application of this network to understand how cells can detect and respond appropriately to the same stimulus (insulin) occurring with different dynamics (sustained low concentration or transient high concentrations). Not only does this study indicate how the liver interprets the fed and fasted states by properly interpreting and decoding the concentration and temporal dynamics of insulin, but this type of trans-omic analysis can be applied to other complex dynamic regulatory systems to understand the principles by which cells interpret dynamic stimuli.

## RESULTS

### Construction of a Trans-omic Network of Insulin Action from Phosphoproteomic, Transcriptomic, and Metabolomic Data

Here, we constructed trans-omic network of insulin action in rat FAO hepatoma cells by integrating phosphoproteomic, transcriptomic, and metabolomic data (Figure 1). We previously measured the phosphoproteome (Yugi et al., 2014) and transcriptome (Sano et al., 2016) in insulin-stimulated FAO cells; however, only a subset of the data were examined in those studies. In this study, we used all phosphoproteome and transcriptome data. Additionally, we measured metabolomic data with concentrations of insulin (0.01, 0.1, 0.3, 1, 3, 10, and 100 nM) for a time course spanning 240 min (Figure 1; Table S6, see STAR Methods). Using these three data sets and information about the enzymatic and allosteric regulation of metabolic enzymes and about activations and inhibitions of TFs, we constructed a trans-omic network of insulin action in FAO cells that included protein phosphorylation, gene expression, metabolites, and allosteric regulation of enzymes by metabolites. Each network was constructed of several layers and these networks were integrated through four steps to produce the trans-omic regulatory network (Figure 1): Step I, construction of the “cellular functions regulatory network” representing phosphorylation-dependent regulation of cellular functions obtained by integration of the phosphoproteomic data with pathways in the Kyoto Encyclopedia of Genes and Genomes (KEGG) database (Kanehisa et al., 2012, 2017); Step II, construction of the “transcriptional regulatory network” containing the TFs predicted as regulators of the insulin-responsive genes (IRGs) obtained from the transcriptomic and connected to insulin signaling; Step III, construction of the “metabolism regulatory network” representing the regulation of metabolic enzymes by phosphorylation and allosteric events obtained from the metabolomic and phosphoproteomic data and allosteric regulatory information in the BRENDA database (Schomburg et al., 2013); Step IV, construction of the trans-omic regulatory network by integrating the three networks from Steps I to III. We further identified how different concentrations and temporal patterns of insulin triggered specific patterns of activity within the trans-omic network (selective regulatory networks) by calculating the sensitivity and time constant for the response to insulin stimulation.

**Figure 1.**
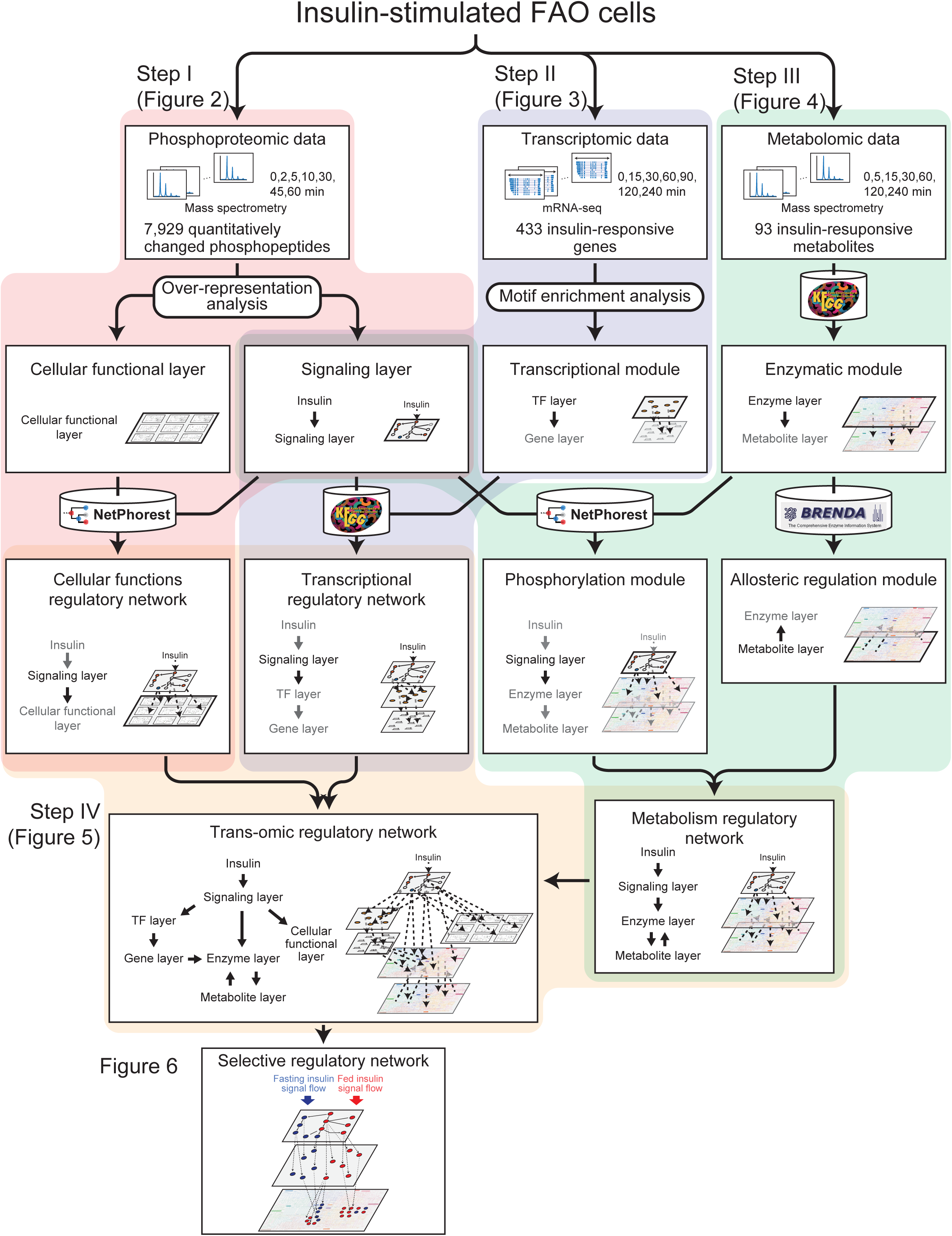
Summary of Procedures for Trans-Omic Network Construction. The trans-omic network was constructed in four main steps by integrating six layers of data into three networks based on phosphoproteome, transcriptome, and metabolome data and defining regulatory connections among the layers. Dark gray labeling of a layer indicates the layer newly identified in the frame; light gray indicates the layer identified in the previous frame. The detailed procedures can be found in STAR Methods.

### Step I: Construction of the “cellular functions regulatory network” obtained by integration of the phosphoproteomic data with pathways in the KEGG database

Previously, we obtained phosphoproteomic data of acute insulin action (< 60 minutes). We used the 199 phosphopeptides from the 49 metabolic enzymes in our previous work (Yugi et al., 2014). For this analysis, we use all 7,929 phosphopeptides from 3,468 proteins. The phosphopeptides included 6,989 phosphoserine (pSer), 1,421 phosphothreonine (pThr), and 79 phosphotyrosine (pTyr) (Figure S1A). Singly phosphorylated peptides represented 93% of the 7,929 phosphopeptides (Figure S1A). We defined insulin-dependent protein phosphorylation as any phosphopeptide exhibiting a change in phosphorylation intensity greater than a 1.5-fold increase or less than a 0.67-fold decrease at one or more time points in response to insulin stimulation. We obtained 3,288 phosphopeptides that changed in response to insulin stimulation; these sites were present on 1,947 proteins. Hereafter, we define a protein with at least one quantitatively changed phosphopeptide as an insulin-responsive phosphoprotein (IRpP). Based on the 1,947 IRpPs, we constructed the cellular functions regulatory network in three steps (Figure 1): Step I-i, identification of the “signaling layer” using IRpPs that are over-represented in KEGG signaling pathways; Step I-ii, identification of the “cellular functional layer” from the selected set of pathways in KEGG in that IRpPs are over-represented; and Step I-iii, connection of “responsible protein kinases” in the signaling layer to IRpPs in the cellular functional layer using the NetPhorest, a kinase-substrate relationship prediction tool.

#### Step I-i: Identification of the “signaling layer”

To generate the signaling layer, we constructed an insulin-dependent phosphorylation of signaling network by integrating all signaling pathways in the KEGG database in which IRpPs were significantly over-represented (Figure S1B; Table S1). Proteins with transcripts that were not expressed in FAO hepatoma cells and proteins that are not located in downstream of InsR in the KEGG database were then removed to define the signaling layer (Figure 2A, see STAR Methods). The signaling layer consisted of 146 proteins and 19 compounds including PIP3, cAMP, and IP3. The proteins included the well-known insulin-responsive kinases Akt, Erk1, Erk2, and mTORC1 (Lizcano and Alessi, 2002; Lu et al., 2012; Saltiel and Kahn, 2001). The signaling layer included 289 phosphopeptides from 125 proteins; 134 of the phosphopeptides quantitatively changed (56 increased, 43 decreased, and 33 both increased and decreased, depending on the time point analyzed).

**Figure 2.**
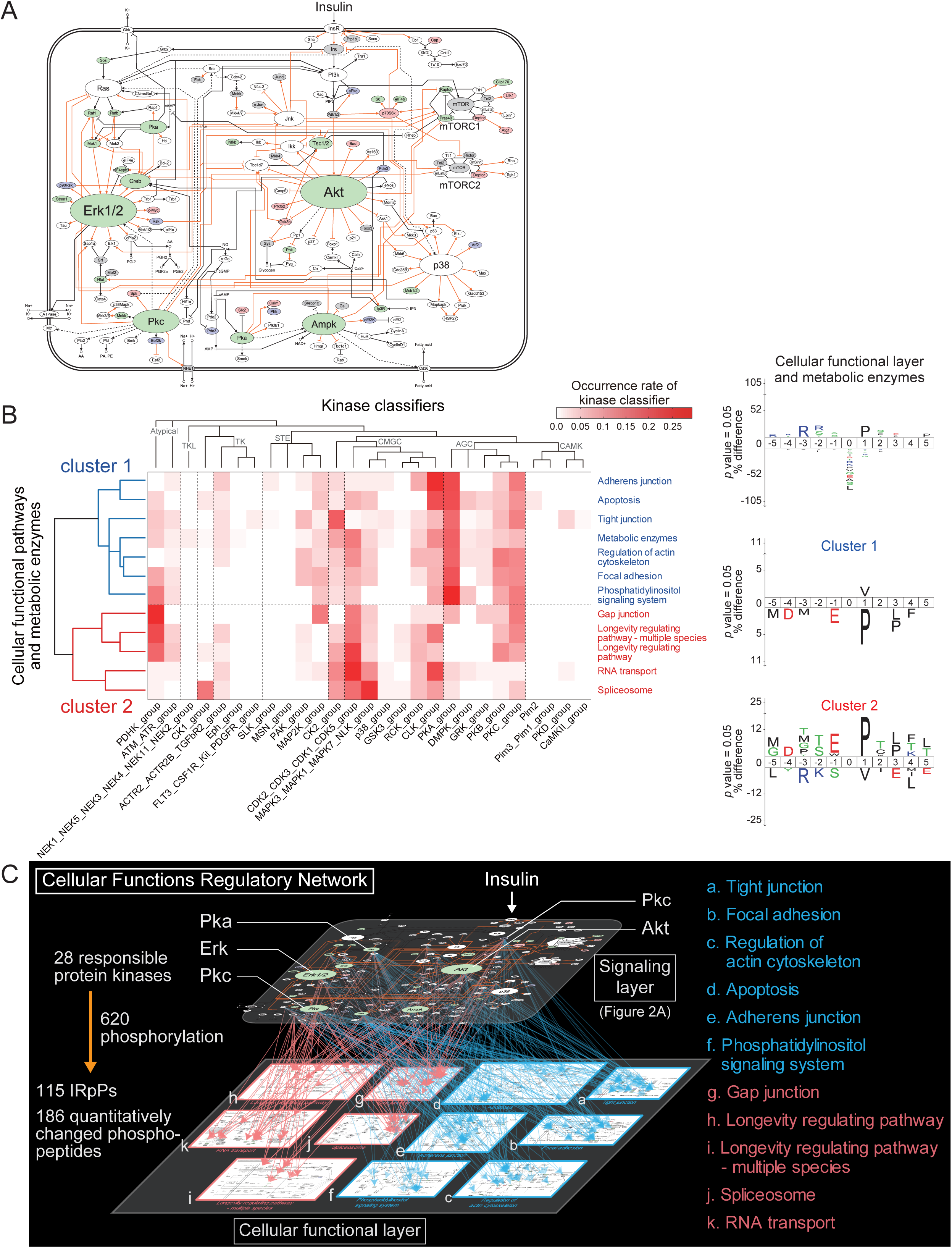
Step I: Construction of the Cellular Functions Regulatory Network from the Phosphoproteomic Data. (A) Signaling layer. Red nodes indicate proteins with an increase (> 1.5-fold) in one or more phosphorylation sites, blue indicates proteins with a decrease (< 0.67-fold), and green nodes indicate proteins with both increased and decreased phosphorylation sites. Gray nodes indicate proteins with unchanged phosphorylation and white indicates proteins with no detected phosphorylation detected. Node size indicates the number of its interactions with other molecules. Orange edges indicate phosphorylation. (B) Clustering of cellular functional pathways and metabolic enzymes by kinase class (left). Amino acid motifs for the quantitatively changed phosphopeptides included in cellular functional layer and metabolic enzymes (right, top), cluster 1 (right, middle), and cluster 2 (right, bottom) are shown as logo plots. The characters above and below each horizontal line indicate amino acids showing significant over-representations and under-representations (p < 0.05), respectively. The height of the letter representing an amino acid at each position reflects the difference in the frequency of its occurrence in the sets of phosphopeptides and reference proteins. (C) Cellular functions regulatory network. The arrows (from the top to the bottom layer) indicate phosphorylation of the quantitatively changed phosphopeptides by the responsible proteins kinases. See also Figure S1 and Table S1, S2, and S3.

#### Step I-ii: Identification of the “cellular functional layer”

To define the cellular functional layer, we collected all KEGG pathways, except for those defined as signaling pathways and those that function specifically in tissues other than liver. We excluded these two groups of pathways, because the signaling pathways are already represented in the previous step and our data are from rat FAO hepatoma cells, which are a liver cell line. From this set of pathways, we extracted those in which IRpPs were significantly over-represented, and defined these 11 pathways as the cellular functional layer (Figure S1B; Table S1).

The cellular functional layer included pathways that regulate cell adhesion, such as *Tight junction* and *Gap junction*; pathways that regulate the cytoskeleton, such as *Regulation of actin cytoskeleton*; and pathways that regulate posttranscriptional processes, such as *RNA transport* and *Spliceosome.* Surprisingly, pathways that regulate metabolism did not have a significant over-representation of IRpPs, possibly because of insufficient detection of phosphopeptides from metabolic enzymes.

#### Step I-iii: Connection of “responsible protein kinases” in the signaling layer to IRpPs in the cellular functional layer

In the final step to generate the cellular functions regulatory network, we used the NetPhorest kinase-substrate recognition prediction tool (Horn et al., 2014; Miller et al., 2008) to assign protein kinases in the signaling layer to the phosphopeptides in the proteins in the cellular functional layer. We defined the responsible protein kinases as those with the highest probability of recognition for each quantitatively changed peptide (see STAR Methods). The cellular functional layer included 1,103 phosphopeptides and 216 IRpPs containing 492 quantitatively changed phosphopeptides. For 216 TRpPs in the cellular functional layer, we predicted 1,688 kinase-substrate relationships between 86 responsible protein kinases and 486 quantitatively changed phosphopeptides in 215 IRpPs in the 11 pathways of cellular functional layer (Table S2).

Next, we identified responsible protein kinases enriched in a pathway by calculating the occurrence rates of responsible protein kinases for the quantitatively changed phosphopeptides present in proteins in the 11 pathways included in the cellular functional layer and in metabolic enzymes from the KEGG database (Table S3, see STAR Methods). We performed clustering analysis with the pathways included in the cellular functional layer and the metabolic enzymes using the occurrence rates of the responsible protein kinases, which resulted in two clusters (Figure 2B, left). Cluster 1 included pathways that regulate cell adhesion and the cytoskeleton, such as *Tight junction, Gap junction,* and *Regulation of actin cytoskeleton;* cluster 2 included pathways that regulate post-transcriptional processes, such as *RNA transport* and *Spliceosome.* We generated the motif logos of amino acid sequences of phosphopeptides in each cluster (Figure 2B, right) (Colaert et al., 2009), and the motif logos of those included in each pathway and those in metabolic enzymes (Figure S1C) (Workman et al., 2005). In quantitatively changed phosphopeptides in proteins in the pathways of the cellular functional layer and in the metabolic enzymes, a proline residue (P) at the +1 position and an arginine residue (R) at the -3 position were significantly over-represented (*p* < 0.05) when compared to the amino acid composition of rat proteins (Figure 2B, right). In cluster 1, only a valine residue (V) at the +1 position was over-represented and P at the +1 position was significantly under-represented in the quantitative changed phosphopeptides. In cluster 2, the P at the +1 position was significantly over-represented in the quantitatively changed phosphopeptides (Figure 2B, right). This result indicates the selective phosphorylation of proteins involved in distinct cellular functions by two different classes of kinases: Basophilic kinases, such as Pka, of the AGC super family of kinases target substrates in cluster 1 and 2, whereas proline-directed kinases, such as the CDKs and MAPKs of the CMGC super family, and PDHKs, of Atypical super family of kinases target substrates in cluster 2 specifically. Of the 86 predicted responsible protein kinases, 28 were part of the signaling layer. The signaling layer and the cellular functional layer were connected by 620 kinase-substrate relationships, represented by the 28 responsible protein kinases in the signaling layer and the 186 quantitatively changed phosphopeptides in 115 IRpPs in the cellular functional layer. These connected layers represent the cellular functions regulatory network (Figure 2C).

### Step II: Construction of the transcriptional regulatory network from transcriptomic and phosphoproteomic data

We previously measured the transcriptome in insulin-stimulated FAO cells and analyzed 13 up-regulated and 16 down-regulated genes in detail (Sano et al., 2016). Here, we used the complete transcriptomic data set, predicted TFs that regulate the IRGs, and connected the proteins in the signaling layer to the TFs to create the a transcriptional regulatory subnetwork in four steps: Step II-i, identification of insulin-responsive genes (IRGs) that are up-regulated or down-regulated in response to insulin; Step II-ii, classification of the up-regulated and down-regulated IRGs according to their sensitivity (*EC*_*50*_) to insulin and the time constant (*T*_*1/2*_) of their response to insulin; Step II-iii, prediction of TFs specific to each class of IRGs; and Step II-iv, connection of the TFs to the signaling layer.

#### Step II-i: identification of IRGs that are up-regulated or down-regulated in response to insulin

Previously, we identified 490 differentially expressed transcripts (Sano et al., 2016) using Cuffdiff (Trapnell et al., 2009, 2012). Among the genes corresponding to these differentially expressed transcripts, we defined 433 genes with fragments per kilobase of transcript per million mapped reads (FPKM) at all time points as insulin-responsive genes (IRGs). We categorized the IRGs into 114 up-regulated, 144 down-regulated, and 175 other according to criteria that identify IRGs with smaller variation and larger responses (Figure S2A-C; Table S4, see STAR Methods).

#### Step II-ii: Classification of the up-regulated and down-regulated IRGs according to sensitivity and time constant

To classify the IRGs, we estimated the sensitivity and the response time of the expression of the IRGs to insulin (see STAR Methods). We estimated sensitivity from the *EC*_*50*_, which we defined as the concentration of insulin that produces 50% of the maximal area under the curve (AUC) of a time series of gene expression (Figure S2D, see STAR Methods). A smaller *EC*_*50*_ indicates a higher sensitivity to insulin (responds at a lower concentration), and vice versa. We estimated response time from the *T*_*1/2*_, which we defined as the time when the change in gene expression reaches 50% of the peak amplitude (Figure S2D, see STAR Methods). A smaller *T*_*1/2*_ indicates a faster response to insulin and vice versa. We calculated the *EC*_*50*_ and *T*_*1/2*_ for each of the up-regulated and down-regulated IRGs. The distributions of the *EC*_*50*_ values and the *T*_*1/2*_ values between the up-regulated and down-regulated IRGs were significantly different (adjusted *p* < 0.01). The average *EC*_*50*_ of the up-regulated IRGs was larger than that of the down-regulated IRGs, and the average *T*_*1/2*_ of the up-regulated IRGs was smaller than that of the down-regulated IRGs (Table 1), consistent with our previous observations (Sano et al., 2016). Furthermore, the distributions of the *EC*_*50*_ and *T*_*1/2*_ values of the up-regulated and down-regulated IRGs showed bimodal distributions (Figure S2E; Table 1), suggesting that the IRGs can be divided into two functional blocks by sensitivity and two functional blocks by time constant; four functional blocs in total. We divided the IRGs into classes on the basis of the *EC*_*50*_ and *T*_*1/2*_ values using Otsu’s thresholding method (Otsu, 1979). This method resulted in four classes: Class 1, high sensitivity (*EC*_*50*_ < threshold) and fast response (*T*_*1/2*_ < threshold); Class 2, high sensitivity and slow response (*T*_*1/2*_ > threshold); Class 3, low sensitivity (*EC*_*50*_ > threshold) and fast response; and Class 4, low sensitivity and slow response (Figure 3A, S2G, and S2H; Table S4). Approximately 50% of the up-regulated IRGs (57 out of 114 up-regulated IRGs) belong to Class 3 (low sensitivity and fast response), and a similar proportion of the down-regulated IRGs (70 out of 144 down-regulated IRGs) belong to Class1 (high sensitivity and fast response) (Figure 3A and B; Table S4). The classified up-regulated and down-regulated IRGs were defined as the “gene layer.”

**Table 1.** Averages and Medians of *EC*_*50*_ and *T*_*1/2*_ values in Insulin-Responsive Genes (IRGs) and Insulin-Responsive Metabolites (IRMs)

| | | Average | Median | $p$ value | Adjusted $p$ value | Mode | |
| --- | --- | --- | --- | --- | --- | --- | --- |
|  |  |  |  |  |  | < threshold | > threshold |
| $EC_{50}$<br>(nM) | Up-regulated IRGs | 1.55 | 2.19 | $9.64 \times 10^{-15}$ | $3.86 \times 10^{-14}$ | 0.25 | 6.31 |
|  | Down-regulated IRGs | 0.25 | 0.16 |  |  | 0.10 | 1.00 |
| | Increased IRMs | 4.90 | 6.61 | $1.34 \times 10^{-10}$ | $5.36 \times 10^{-10}$ | 0.40 | 1.58 |
|  | Decreased IRMs | 1.29 | 1.23 |  |  | 0.06 | 0.63 |
| $T_{1/2}$<br>(min) | Up-regulated IRGs | 67.56 | 50.15 | $9.82 \times 10^{-5}$ | $3.93 \times 10^{-4}$ | 45.00 | 120.00 |
|  | Down-regulated IRGs | 86.86 | 58.51 |  |  | 45.00 | 120.00 |
| | Increased IRMs | 56.14 | 67.93 | $1.90 \times 10^{-3}$ | $7.60 \times 10^{-3}$ | 0.00 | 105.00 |
|  | Decreased IRMs | 17.97 | 10.93 |  |  | 15.00 | 60.00 |

**Figure 3.**
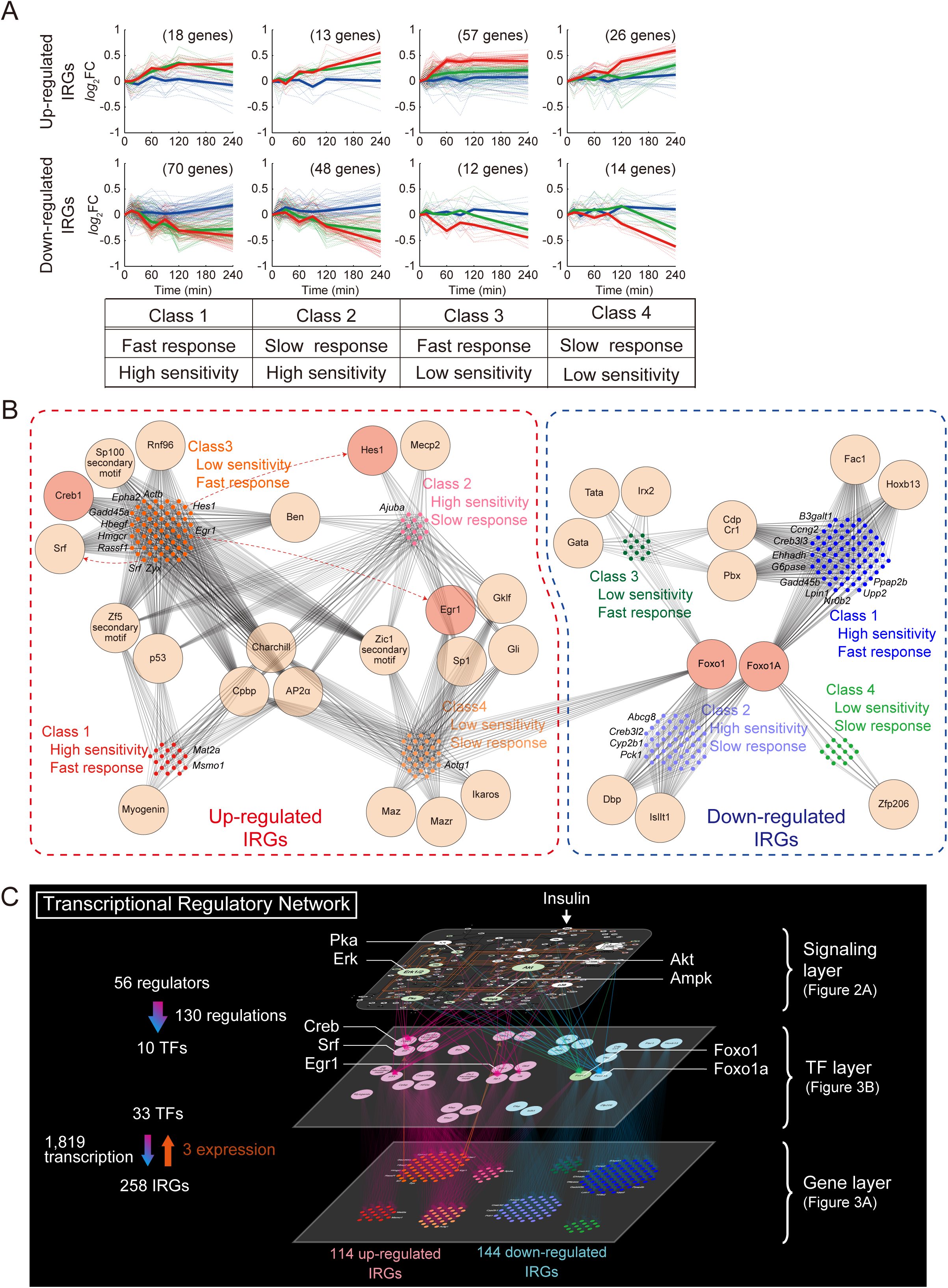
Step II: Construction of the Transcriptional Regulatory Network from the Transcriptomic and the Phosphoproteomic Data. (A) Time courses of IRGs in each class with high or low sensitivity and with fast or slow time constant. Dashed lines indicate the time series of each IRG. The y-axis indicates the base 2 logarithm of fold change against time 0 (*log*_2_FC). Blue, green, and red lines indicate responses to 0.01, 1, and 100 nM insulin, respectively, with bold lines representing the averaged responses at each concentration. (B) Transcriptional module. Dots indicated IRGs, circles indicate TFs, gray lines indicate transcriptional regulation, and red dashed arrows connect a TF encoding gene to the product TF. Dark orange TFs indicate TFs measured for phosphorylation or protein abundance by Western blotting. (C) Transcriptional regulatory network. See also Figure S2, S4, and Table S4 and S5.

#### Step II-iii: Prediction of TFs specific to each class of IRGs

To assign TFs to each of the IRGs, we identified putative binding motifs for TFs in the flanking region [-300 bp to +100 bp of the consensus transcription start site (Arner et al., 2015; Kinsella et al., 2011)] of each IRG using TRANSFAC (Matys et al., 2006) and Match (Kel et al., 2003). Subsequently, we used motif enrichment analysis of the TF binding motifs to assign TFs to each class of up-regulated and down-regulated IRGs (see STAR Methods).

For the 433 IRGs, we identified 168 putative TF binding motifs. With the motif enrichment analysis we identified 33 TFs as likely regulators of the up-regulated and the down-regulated IRGs (Table S5). The 33 TFs mapped to four classes of IRGs were defined as the “TF layer,” and the relationship between the 33 TFs and the 114 up-regulated and the 144 down-regulated IRGs was denoted as the “transcriptional module” (Figure 1, Figure 3B). Twenty-two TFs were assigned to the classes of up-regulated IRGs and 12 TFs to the classes of down-regulated IRGs. The TFs assigned to the up-regulated or down-regulated IRGs were mutually exclusive except for Foxo1. Foxo1 was connected to IRGs in all classes of down-regulated IRGs and in up-regulated IRGs in Class 4.

#### Step II-iv: Connection of the TFs to the signaling layer

To connect the signaling layer to the 33 TFs predicted to regulate the IRGs, we used the pathway information in the KEGG database. We identified 117 regulators related to 12 of the TFs. The 56 of the 117 regulators were present in the signaling layer, and connected to 10 of the IRG-regulating TFs, resulting in the transcriptional regulatory network (Figure 3C). This transcriptional regulatory network indicates that the Erk pathway was a primary regulator of the up-regulated IRGs, because the TFs associated with these IRGs include Egr1, Creb1, and Srf. Consistent with this inference, these TFs are known targets of the Erk MAPK pathway (Deak et al., 1998; Murphy et al., 2004; Nakayama et al., 2008; Shaul and Seger, 2007). In contrast, Foxo1, which was connected to all four classes of the down-regulated IRGs, was connected to Akt in the signaling layer, which is consistent with the known regulation of Foxo1 by the Akt pathway (Barthel et al., 2005; Matsuzaki et al., 2003; Mounier and Posner, 2006; Puigserver et al., 2003). These results validate the accuracy of the transcriptional regulatory network.

### Step III: Construction of the metabolism regulatory network from metabolomic and phosphoproteomic data

We obtained metabolomic data with nine concentrations of insulin over a time course of 240 min and used these data, along with phosphoproteomic data obtained under a single concentration (1 nM) of insulin for a period of 60 minutes, to construct a phosphorylation-dependent metabolism regulatory network according to our previous method (Yugi et al., 2014). We assumed that many of protein phosphorylation changes induced by insulin occur within 60 min (Figure S4A), whereas we assume that subsequent changes in gene expression and metabolic activity would continue beyond this time. We integrated the metabolomic data with the phosphoproteomic data through a layer of metabolic enzymes responsible for the changes in the metabolites in three steps: Step III-i, identification of insulin-responsive metabolites (IRMs) and their responsible metabolic enzymes; Step III-ii, identification of allosteric regulation of responsible metabolic enzymes by IRMs; and Step III-iii, connection of responsible protein kinases in the signaling layer to the quantitatively changed phosphopeptides on responsible metabolic enzymes by predicting kinase-substrate relationships.

#### Step III-i: Identification of IRMs and their responsible metabolic enzymes

To identify a high confidence set of IRMs from multiple experimental data sets, we performed a three-way analysis of variance (ANOVA) with the insulin doses, stimulation times, and data acquired on different days as main factors (see STAR Methods). This process identified 93 IRMs that showed significant changes (*FDR* < 0.1) in response to insulin stimulation (Table S6). As with IRGs, we categorized the 93 IRMs into 42 increased, 43 decreased, and eight other IRMs according to criteria that identify IRMs with larger responses (Figure S3A; Table S6, see STAR Methods). We used only those that consistently increased or decreased for construction of the network; we did not use the IRMs in the other category further. In the central carbon metabolism (glycolysis and gluconeogenesis, the TCA cycle, and the pentose phosphate pathway), multiple metabolites exhibited insulin-induced changes in abundance (Figure 4A and B). Glucose 6-phosphate (G6P) and fructose 6-phosphate (F6P) were IRMs that decreased. In contrast, metabolites downstream of F6P in glycolysis—fructose 1,6-biphosphate (F1,6BP), 3-phospho-D-glyceroyl phosphate (3PG), and phosphoenolpyruvate (PEP)—were increased. Several metabolites in the TCA cycle (citrate, 2-oxoglutarate, succinate, fumarate, and malate) were also increased in response to insulin. Although many amino acids decreased, Ala, Arg, and Ser increased (Figure 4A).

**Figure 4.**
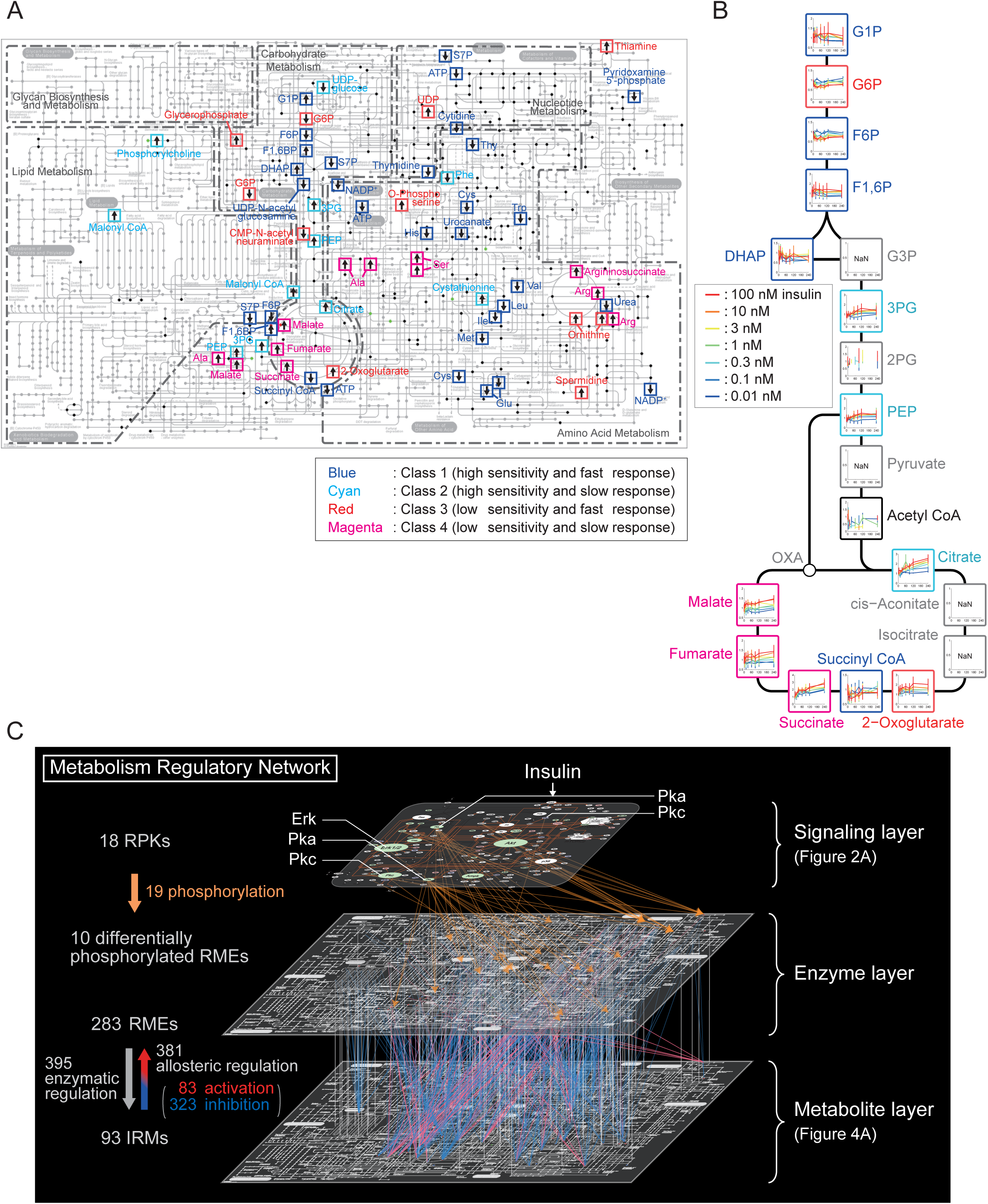
Step III: Construction of the Metabolism Regulatory Network from the Metabolomic and Phosphoproteomic Data. (A) IRMs projected on the KEGG *metabolic pathways.* Arrows indicate whether an IRM increased or decreased. The colors of the box outlines indicate the classes of the IRMs. (B) Metabolites in the central carbon metabolism. The black frames indicate the metabolites that did not show significant changes in response to insulin in a three-way ANOVA, and gray frames indicate those with unmeasured points at one and more time points (NaN). The other colors correspond to the classes. (C) Metabolism regulatory network. RMEs, responsible metabolic enzymes; RPKs, responsible protein kinases. See also Figure S3 and Table S6, S7, S8, and S9.

Similar to the analysis of IRGs, to estimate the sensitivity and response time of the IRMs to insulin, we calculated the *EC*_*50*_ and *T*_*1/2*_ values (Figure S2D and S3A, B, Table 1). We defined *EC*_*50*_ as the concentration of insulin that produces 50% of the maximal AUC of a time series of gene expression (Figure S2D, see STAR Methods) and *T*_*1/2*_ as the time when the change in gene expression reaches 50% of the peak amplitude (Figure S2D, see STAR Methods). The distributions of the *EC*_*50*_ values and the *T*_*1/2*_ values were significantly different (adjusted *p* < 0.01) between the increased and decreased IRMs, and the average values of the *EC*_*50*_ and the *T*_*1/2*_ for the increased IRMs were larger than those for the decreased IRMs (Table 1). Thus, on average IRMs that decreased responded more quickly and with higher sensitivity than did the IRMs that increased in response to insulin.

Furthermore, the distributions of *EC*_*50*_ values of both the increased and decreased IRMs were unimodal (Figure S3B). The distribution of the *T*_*1/2*_ of the increased IRMs was bimodal; that of the decreased IRMs was unimodal (Table 1, Figure S3B). Using Otsu’s method (Otsu, 1979), we determined the thresholds of the *EC*_*50*_ and *T*_*1/2*_ values and then classified the IRMs into four classes analogous to the classes for the IRGs: Class 1, high sensitivity and fast response and; Class 2, high sensitivity and slow response; Class 3, low sensitivity and fast response, and Class 4, low sensitivity and slow response (Figure 4A and B). We mapped the four classes of IRMs onto the KEGG *metabolic pathways* to produce the “metabolite layer.”

The abundances of the metabolites change depending on the reaction rates of the synthesis (influx) and the consumption (efflux) of the metabolite itself, these rates are determined by activities of metabolic enzymes, the amounts of the enzymes, and the amounts of the substrates and products. We defined the metabolic enzymes that directly produce or consume at least one IRM as responsible metabolic enzymes. Using the KEGG database, we identified 183 and 115 responsible metabolic enzymes for 50 IRMs as substrate and for 39 IRMs as product (Figure S3), respectively. The responsible metabolic enzymes identified for the IRMs as substrates and for those as products were not mutually exclusive. Moreover, the IRMs identified as substrates and those as products were not mutually exclusive. We identified total of 283 responsible metabolic enzymes for total 62 of the 93 IRMs (Figure S3; Table S7). We mapped the responsible metabolic enzymes onto the KEGG *metabolic pathways* to produce the “enzyme layer,” and the relationship between the 283 responsible metabolic enzymes and the 62 IRMs was denoted as the “enzymatic module” (Figure 1).

#### Step III-ii: Identification of allosteric regulation of responsible metabolic enzymes by IRMs

Many metabolic enzymes are regulated allosterically by metabolites; therefore, we identified IRMs that function as allosteric regulators for the responsible metabolic enzymes using the BRENDA database (see STAR Methods), which is a database with information regarding allosteric effectors and their target enzymes (Schomburg et al., 2013). A metabolite can operate as an activator for some enzymes and as an inhibitor for others. We found 70 IRMs that function as allosteric effectors for 146 of the responsible metabolic enzymes. These interactions produced 381 allosteric regulatory events (83 activating events and 323 inhibitory events) (Figure S3; Table S8). To generate the “allosteric regulation module,” we connected these allosteric-regulating IRMs to the responsible metabolic enzymes through the 381 allosteric regulatory events (Figure 1).

#### Step III-iii: Connection of responsible protein kinases in the signaling layer to the quantitatively changed phosphopeptides on responsible metabolic enzymes

Metabolic enzyme activity can also be regulated through changes in phosphorylation. Therefore, we examined the phosphoproteomic data for insulin-responsive changes in the phosphorylation of the responsible metabolic enzymes. We identified 174 phosphopeptides from 51 responsible metabolic enzymes. Only 77 quantitatively changed phosphopeptides on 30 enzymes exhibited insulin-stimulated changes in abundance (Table S7). These 30 metabolic enzymes corresponded to 29 IRMs (Table S7). Similar to how we connected the signaling layer to the cellular functional layer to create the cellular functions regulatory network, we used kinase-substrate relationship analysis and probability analysis in NetPhorest to connect kinases in signaling layer to the quantitatively changed phosphopeptides in metabolic enzymes (Figure 4C and see STAR Methods). We predicted 206 kinase-substrate relationships between 49 responsible protein kinases and 77 quantitatively changed phosphopeptides in 30 responsible metabolic enzymes.

Of the 49 responsible protein kinases, 18 were present in the signaling layer, and these 18 kinases were associated with 20 quantitatively changed phosphopeptides in 10 responsible metabolic enzymes through 19 kinase-substrate relationships (Figure 4C). Thus, the metabolism regulatory network contained the 19 phosphorylation events mediated by the 18 responsible protein kinases on the 10 responsible metabolic enzymes, and the 381 allosteric regulatory events (83 activating events and 323 inhibitory events) between 70 IRMs and 146 responsible metabolic enzymes (Figure 4C).

### Step IV: Construction of the trans-omic regulatory network

To construct the trans-omic regulatory network (Figure 5), we integrated the cellular functions regulatory network (Figure 2C, Step I), the transcriptional regulatory network (Figure 3C, Step II), and the metabolism regulatory network (Figure 4C, Step III). The responsible metabolic enzymes that were involved in IRpPs include rate-limiting enzymes of glycolysis and gluconeogenesis, 6-phosphofructo-2-kinase/fructose-2,6-biphosphatase 1 (Pfkfb) and liver type phosphofructokinase (Pfkl), and rate-limiting enzymes of fatty acid synthesis, ATP citrate lyase (Acly), acetyl-CoA carboxylase alpha (Acaca), and fatty acid synthase (Fasn). Responsible metabolic enzymes involved in the down-regulated IRGs were G6pase, Pck1, pyruvate kinase, liver and RBC (Pklr), and patatin-like phospholipase domain containing 6 (Pnpla6). A responsible metabolic enzyme involved in the up-regulated IRG was methionine adenosyltransferase 2A (Mat2a). *G6Pase* and *Pck1* encode rate-limiting enzymes of gluconeogenesis. Among 93 IRMs, 70 were identified as allosteric effectors. Allosteric regulatory events far exceeded the number of phosphorylation-mediated or transcriptional regulatory events for metabolic enzymes. This disparity suggested that insulin acts through protein phosphorylation and gene expression to control a few key regulatory points (rate-limiting enzymes) in cellular metabolism. As cellular metabolism changes, numerous metabolites in multiple metabolic pathways change, which thereby changes the abundance and balance of many allosteric effectors. These allosteric effectors function as secondary regulators of insulin to propagate the signal and influence the activity of more metabolic enzymes than those initially regulated by phosphorylation or transcriptional events.

**Figure 5.**
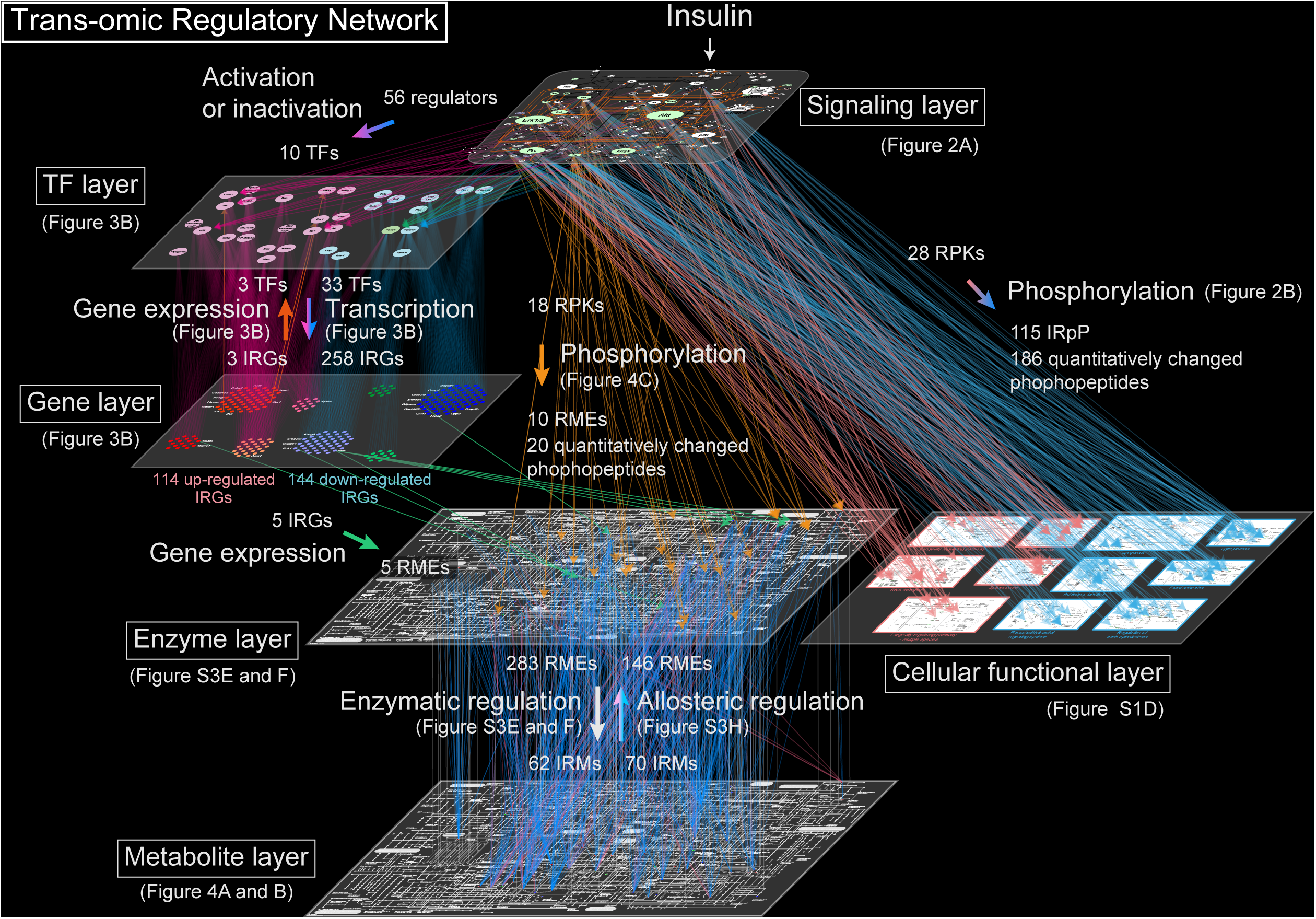
Step IV: Construction of the Trans-Omic Regulatory Network. The trans-omic regulatory network contains six layers and the regulatory relationships among them. It was constructed by integrating the cellular functions regulatory network (Figure 2C, Step I), the transcriptional regulatory network (Figure 3C, Step II), and the metabolism regulatory network (Figure 4C, Step III). RMEs, responsible metabolic enzymes; RPKs, responsible protein kinases.

### Selective fasting and fed insulin signal flow in the trans-omic regulatory network

The pancreas releases insulin in specific temporal patterns and various concentrations. Basal secretion during fasting results in a low circulating concentration with a sustained temporal pattern. Glucose-stimulated secretion in response to meals results in high concentrations of circulating insulin with rapid, transient kinetics (Polonsky et al., 1988) (Figure 6E). To identify selective information flow through the trans-omic regulatory network in response to the concentration and temporal pattern of insulin, we used the *EC*_*50*_ and *T*_*1/2*_ values for molecules in the signaling layer, the TFs, the IRGs, and IRMs. We could calculate these values for the IRGs and IRMs from the transcriptomic and metabolomic data, because those data were acquired from multiple conditions of insulin stimulation. In contrast, we could not calculate these values for protein phosphorylation or protein abundance from the phosphoproteomic data, because those data were obtained from only a single insulin stimulus. Therefore, we selected key signaling molecules and performed insulin dose-dependent time course analysis data with different concentrations of insulin.

**Figure 6.**
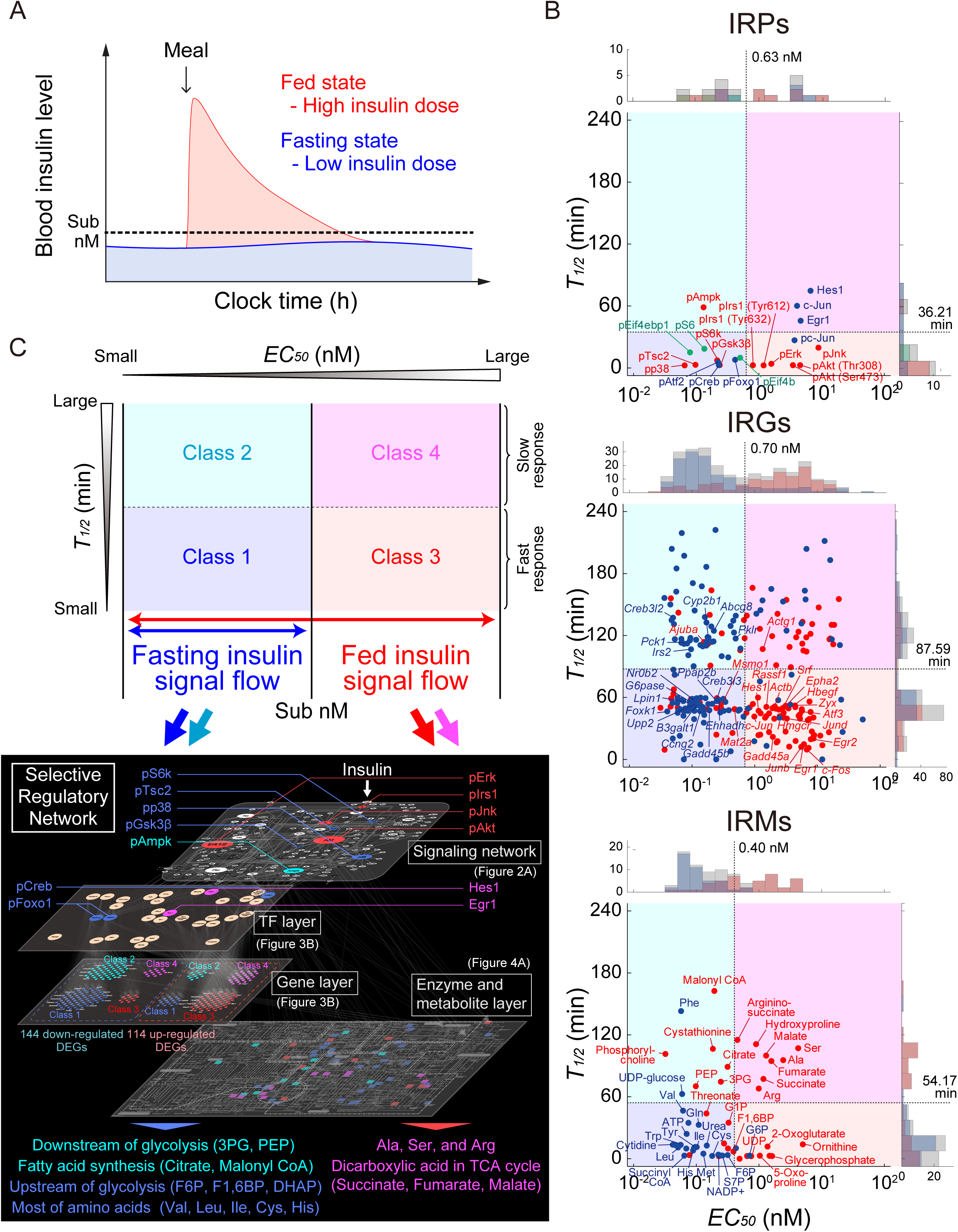
Identification of Fasting and Fed Insulin Signal Flow in the Trans-omic Regulatory Network. (A) *In vivo* temporal patterns of insulin (Polonsky et al., 1988). (B) The distributions of *EC*_*50*_ and *T*_*1/2*_ values of IRPs (top), IRGs (middle), and IRMs (bottom). In the IRP graph, red, blue, and green dots indicate signaling factors, TFs, and protein synthesis-related factors, respectively. In the IRG graph, red dots in the IRG graph indicate up-regulated IRGs, blue dots indicate down-regulated IRGs. In the IRM graph, red dots indicate increased IRMs and blue dots indicated decreased IRMs. The dotted lines indicate the thresholds of the *EC*_*50*_ and *T*_*1/2*_ values. For simplicity, only the proteins, IRGs and IRMs annotated in the trans-omic regulatory network in C are labeled on the graphs. (C) Classification by *EC*_*50*_ and *T*_*1/2*_ values according to the thresholds and color-coding of the trans-omic regulatory network layers by class. Class 1 and 2 are in high sensitivity network that responds to basal insulin secretion (fasting insulin signal flow) (blue and cyan), and Class 3 and 4 are in the low sensitivity network that responds to meal-stimulated insulin secretion (fed insulin signal flow) (red and magenta). See also Figure S4 and Table S4, S6, and S9.

We limited our analysis to proteins for which antibodies were available (Figure S4A and B). We selected nine signaling proteins to quantify insulin-stimulated changes in phosphorylation: Irs1 (pIrs1), Akt (pAkt), S6k (pS6k), Gsk3β (pGsk3β), Erk (pErk), p38 (pp38), Jnk (pJnk), Ampk (pAmpk), and Tsc2 (pTsc2). We chose these because each of these was connected to many other signaling factors in the signaling layer (Figure 2A). We selected seven TFs. We quantified the phosphorylation of Foxo1 (pFoxo1), Creb (pCreb), Atf2 (pAtf2), and c-Jun (pc-Jun), and the abundances of Egr1, c-Jun, and Hes1. We selected three proteins involved in protein synthesis that are regulated by phosphorylation: S6 (pS6), eIf4ebp1 (peIf4ebp1), and eIf4b (peIf4b). We refer to the selected proteins as insulin-regulated proteins (IRPs).

We estimated the *EC*_*50*_ and the *T*_*1/2*_ values of the IRPs (Figure 6A and S4; Table S9). We divided the IRPs into classes by determining thresholds of the *EC*_*50*_ and *T*_*1/2*_ values using Otsu’s method (Otsu, 1979). We classified IRPs, as we did the IRGs and IRMs, into four classes: Class 1, high sensitivity and fast response and; Class 2, high sensitivity and slow response; Class 3, low sensitivity and fast response, and Class 4, low sensitivity and slow response (Figure 6A; Table S9). The similarity among the thresholds of the *EC*_*50*_ values for IRPs (0.63 nM), IRGs (0.70 nM), and IRMs (0.40 nM) (Figure 6A-C) indicated each of these has subsets that respond differentially to fasting or fed insulin stimuli, because basal secretion in the fasting state is tens to hundreds of pM insulin and meal-induced secretion results in concentrations of insulin in the nM range (Figure 6E, left) (Lindsay et al., 2003; Polonsky et al., 1988). The threshold of the *T*_*1/2*_ for IRPs was fastest at 36.21 min with changes in IRMs occurring next (54.17 min) and changes in IRGs occurring latest (87.59 min) (Figure 6A-C). This is consistent with a delay between the IRPs, which receive the insulin stimulus, and changes in metabolism and gene expression.

Using this information, we identified the components in the four classes of IRPs, IRGs, and IRMs from the trans-omic regulatory network and constructed a selective regulatory network for each class (Figure 6D and E). Perhaps not surprisingly, phosphorylation-mediated regulatory events occurred in Classes 1 and 3, which exhibit a fast response to either low or stimulated insulin concentrations. These changes were transmitted to a subset of metabolites, including those upstream of F6P in glycolysis and most amino acids, which fell in Class 1. Other metabolites changed with a slower time course with some changing in response to low concentrations of insulin and some responding to high concentrations of insulin (Figure 6E).

## DISCUSSION

We showed selective information flow through the trans-omic regulatory network according to the concentration and time of insulin stimulation (Figure 6). We denoted high sensitivity networks as representing the response to basal insulin secretion (fasting insulin signal flow) (Figure 6E, blue and cyan) and low sensitivity networks as representing the response to meal-stimulated insulin secretion (fed insulin signal flow) (Figure 6D and E, red and magenta). In the signaling layer of the trans-omic regulatory network, pIrs1, pJnk, pAkt, and pErk show low sensitivity and fast response (Class 3); thus, these proteins function in the low sensitivity network (Figure 6A, 6E, and S4; Table S9). As such, these proteins can respond to a wide dynamic range of insulin secretion, and therefore can encode both fasting and fed insulin signals. In contrast, pS6k, pTsc2, pp38, and pGsk3β show high sensitivity and fast response; thus, these proteins function in the high sensitivity network (Figure 6A, 6E, and S4; Table S9). This set of proteins respond to low concentrations of insulin and likely to selectively respond to basal insulin secretion.

For TFs, pCreb and pFoxo1 (Class 1) were in the high sensitivity network, whereas Egr1 and Hes1 (Class 4) were part of the low sensitivity network (Figure 6A, 6E, and S4; Table S9). The TFs also segregated into those that responded quickly (pCreb and pFoxo1), which were regulated by phosphorylation by kinases in the signaling layer, such as Erk and Akt (Figure 3C and 5) (Barthel et al., 2005; Deak et al., 1998; Lu et al., 2012; Matsuzaki et al., 2003). In contrast, Egr1 and Hes1 responded slowly, consistent with their regulation by gene expression (Figure 3B, 3C and 5, Table S9) (Herzig et al., 2003; Lemke et al., 2008; Sukhatme et al., 1988; Thiel and Cibelli, 2002).

For the IRGs, most of the down-regulated IRGs (Class 1 and 2) are in the high sensitivity network; most of the up-regulated IRGs (Class 3 and 4) are in the low sensitivity network (Figure 3A, 3B, 6B, 6D and 6E; Table S4). Those in the high sensitivity network include *G6Pase* and *Pck1*, which are the rate-limiting enzymes of gluconeogenesis and are responsive to basal insulin secretion (Lu et al., 2012; Saltiel and Kahn, 2001). *Hmgcr*, which encodes the rate-limiting enzyme for cholesterol synthesis (Durr and Rudney, 1960; Lindgren et al., 1985), is in the low sensitivity network. The identification of these genes in these different pathways is consistent with the physiological roles of the encoded products in the regulation of metabolism by insulin and with our previous studies (Kubota et al., 2012; Noguchi et al., 2013; Sano et al., 2016).

For IRMs in central carbon metabolism, Class 1 metabolites included several that are upstream of glycolysis (F6P, F1,6BP, and DHAP) and Class 2 metabolites included those that are downstream of glycolysis (3PG and PEP) and those involved in fatty acid synthesis (citrate and malonyl CoA) (Figure 4A, 4B, 6C, 6D, and 6E; Table S6). These are the metabolites that would respond to basal insulin secretion as part of the high sensitivity network. Class 1 are the fast-response and Class 2 are the slow-response metabolites, indicating insulin would primarily affect events before 3PG in glycolysis, leading to subsequent changes in the glycolytic pathway and fatty acid synthesis. Class 4 metabolites, such as those that a part of the TCA cycle (succinate, fumarate, and malate), would respond to meal-stimulated insulin secretion as part of the low sensitivity network. These results indicate that the pattern of insulin stimulation divides the central carbon metabolism into three functional blocks: (i) a block upstream of glycolysis, (ii) a block with metabolites that are downstream of glycolysis and tricarboxylic acid in the TCA cycle (citrate), and (iii) dicarboxylic acids in TCA cycle (succinate, fumarate, and malate). Basal insulin secretion in the fasting state engages the first block, then changes in the metabolite of the first block affect the second block. The third block is engaged by insulin secretion in response to eating. For IRMs in amino acid metabolism, most of amino acids such as the branched chain amino acids (Class 1), which were decreased by insulin stimulation, were part of the high sensitivity network; whereas Ala, Ser, and Arg (Class 4), which increased by insulin stimulation and are only a few enzyme steps away from central carbon metabolism, are part of the low sensitivity network that responds to insulin secretion stimulated by eating (Figure 4A, 4B, 6C, 6D, and 6E; Table S6). This result indicates that the pattern of insulin stimulation divides amino acid metabolism into two functional blocks.

We detailed a method for the construction of a trans-omic network by integrating three networks based on phosphoproteome, transcriptome and metabolome data in insulin-stimulated FAO cells. The trans-omic regulatory network contains six layers generated from different types of data, which are then connected through various types of regulatory relationships. We acknowledge that some data and regulatory modules that are needed for a comprehensive trans-omics network are not included in the trans-omic regulatory network. These missing elements include proteome abundance, other types of posttranslational modifications, protein-protein interactions (PPIs), metabolic flux, and epigenomic data. Additionally, some of regulatory relationships in the trans-omic regulatory network are inferred or predicted, which could lead to inaccuracies in the network. For the transcriptional module, we used a limited genomic region surrounding the consensus transcription start site as the flanking region of each IRG to predict connections to TFs. Enhancers and other genomic regions were not considered. Thus, the TF layer may be incomplete. Because many TFs recognize similar consensus motifs of base sequences, some of the regulatory relationships between TFs and IRGs might be inaccurately predicted.

For phosphorylation module, the similarities in phosphorylation site consensus motifs shared by many protein kinases may also have resulted in inaccurate predictions of kinase-substrate relationships. Despite the limitations in the trans-omic regulatory network, we obtained molecular insights into how cells interpret insulin stimulation through the process of creating the various layers necessary to construct the three networks, as well as by evaluation of the trans-omic network.

In this study, we constructed a trans-omic regulatory network from phosphoproteomic, transcriptomic, and metabolomic data, and that incorporates and reveals the selective regulation of pathways through the network in response to different concentrations and duration of insulin stimulus by quantifying sensitivity and time constants for the omic data. Future studies can expand this trans-omic regulatory network for insulin to include other types of data. We propose that these methods of construction of trans-omic regulatory networks and identification of the selective regulatory networks through the trans-omic regulatory network can be applied to explore dynamic cellular responses to other dynamic stimuli. Comparing sensitivities and time constants of molecules in a trans-omic network with *in vivo* concentrations and temporal patterns of extracellular stimuli enables the identification of specific pathways and molecules in a trans-omic regulatory network engaged by different stimulation conditions.

## AUTHOR CONTRIBUTIONS

K. Kawata, K.Y., A.H., and S.K. conceived the project. K. Kawata, K.Y., T.K., Y.T., T. Sano, K.Y.T., M.F., and S.U. analyzed the data. A.H., H.K., and S.K. designed the experiments. A.H. performed the western blotting experiments. M.M. and K.I.N. performed the phosphoproteome measurements. Y.S. performed the RNA-seq experiments. K.S., K. Kato, A.U., M.O., and T. Soga performed the metabolome measurements. K. Kawata, K.Y., A.H., and S.K. wrote the manuscript.

## ACKNOWLEDGEMENTS

We thank our laboratory members for critically reading this manuscript and for their technical assistance with the experiments. The computational analysis of this work was performed in part with support of the super computer system of National Institute of Genetics (NIG), Research Organization of Information and Systems (ROIS). This manuscript was edited by Nancy R. Gough (BioSerendipity, LLC). This work was supported by the Creation of Fundamental Technologies for Understanding and Control of Biosystem Dynamics, CREST (JPMJCR12W3) from the Japan Science and Technology Agency (JST) and by the Japan Society for the Promotion of Science (JSPS) KAKENHI Grant Number (17H06300). K.Y. receives funding from a Grant-in-Aid for Young Scientists (A) (15H05582) from JSPS, and ‘Creation of Innovative Technology for Medical Applications Based on the Global Analyses and Regulation of Disease-Related Metabolites’, PRESTO, from JST. H.K. was supported by JSPS KAKENHI Grant Number 16H06577. K.I.N. and Y.S. were supported by the JSPS KAKENHI Grant Number (17H06301) and (17H06306), respectively. T.S. receives funding from the AMED-CREST from the Japan Agency for Medical Research and Development, AMED.

## SUPPLEMENTAL FIGURE LEGENDS

**Figure S1.**
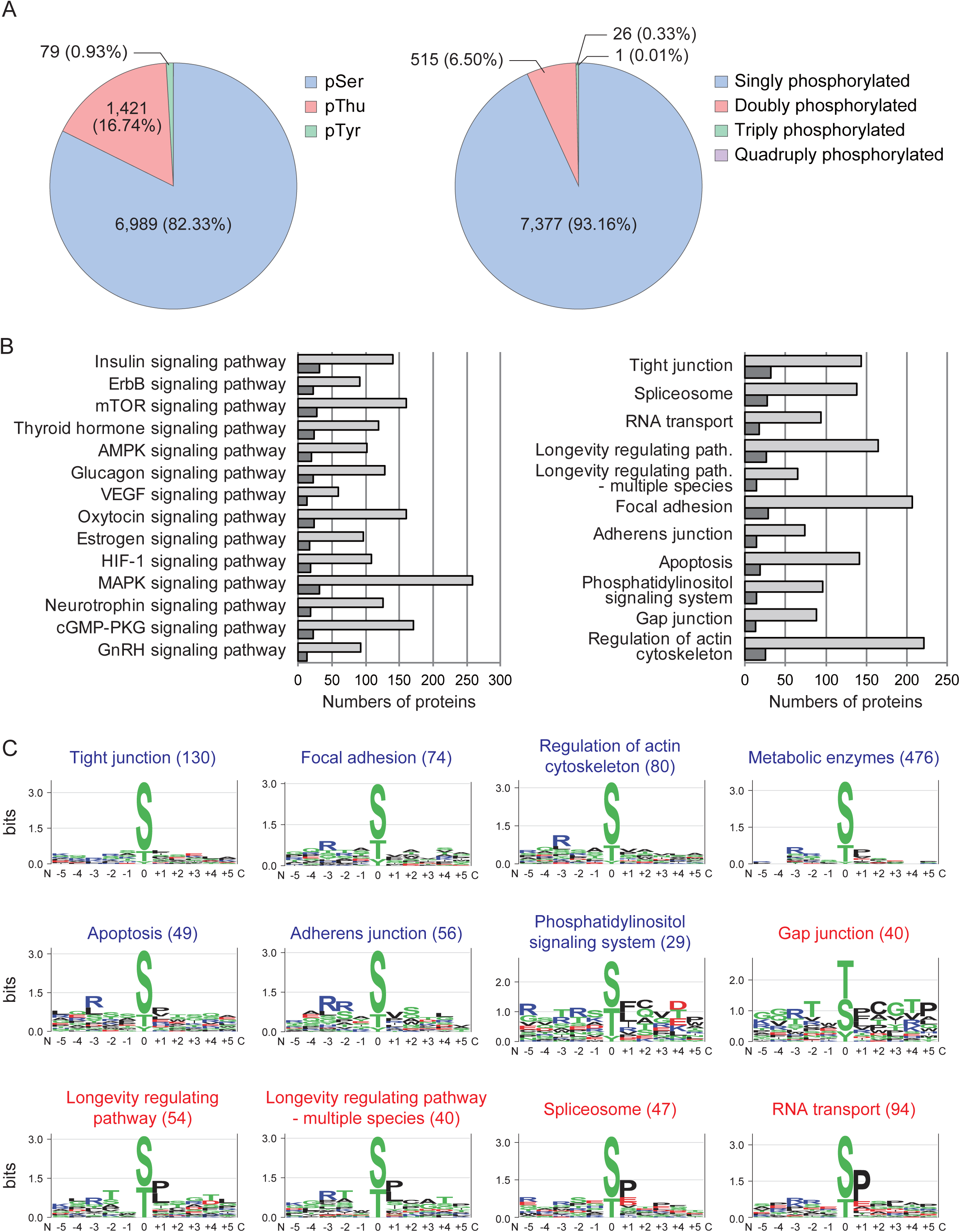
Defining the signaling layer and cellular functional layer for the cellular functions regulatory network, Related to Figure 2. (A) Distribution of phosphorylation sites by amino acid residues and number of phosphorylated sites per peptide. Numbers and parentheses indicate the number of phosphorylation sites or phosphopeptides and percentage relative to the total phosphorylation sites or phosphopeptides, respectively. (B) Number of proteins in the KEGG signaling pathways and the cellular functions pathways. The light gray bars indicate the total numbers of proteins and the dark gray bars indicate the numbers of IRpP contained in each signaling pathway. See also TableS1. (C) Motif logo of amino acid sequences of phosphopeptides in each of the indicated pathways and the metabolic enzymes. The height of each letter at each position is scaled relative to the information content, reflecting the frequency of the corresponding amino acid. Blue pathway names correspond to those in cluster 1; red names correspond to those in cluster 2. The number in parentheses attached to each pathway name represents the number of quantitatively changed peptides included in the pathway.

**Figure S2.**
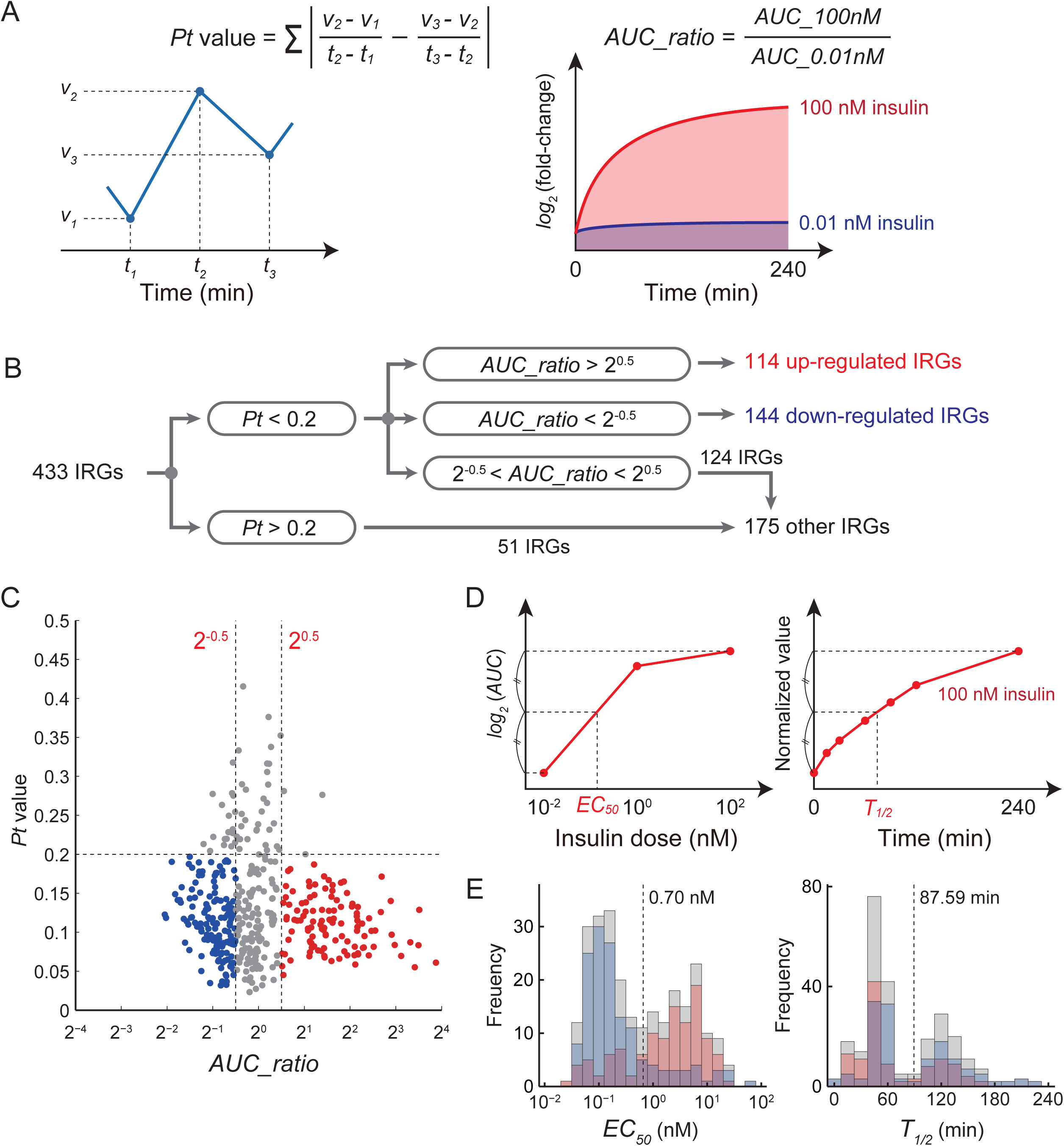
Defining *EC*_*50*_ and *T*_*1/2*_ to classify IRGs, Related to Figure 3. (A) Definition of *Pt* value used an index of expression variation and definition of *AUC_ratio* used as an index of response. (B) Definition of the up-regulated and down-regulated IRGs. (C) Distribution of *Pt* value and *AUC_ratio* in the up-regulated IRGs (red) and down-regulated IRGs (blue). Gray dots indicate the IRGs defined as neither up-regulated nor down-regulated IRGs. Horizontal and vertical dotted lines indicate thresholds of *Pt* values and *AUC_ratios*, respectively. (D) Definition of *EC*_*50*_ used as an index of sensitivity to insulin doses and definition of *T*_*1/2*_ used as an index of time constant. (E) Distribution of *EC*_*50*_ and *T*_*1/2*_ values calculated for up-regulated IRGs (red bars) and down-regulated IRGs (blue bars). Gray bars indicate the sum of the frequency of up-regulated and down-regulated IRGs. The dashed lines indicate the thresholds of the bimodal distributions.

**Figure S3.**
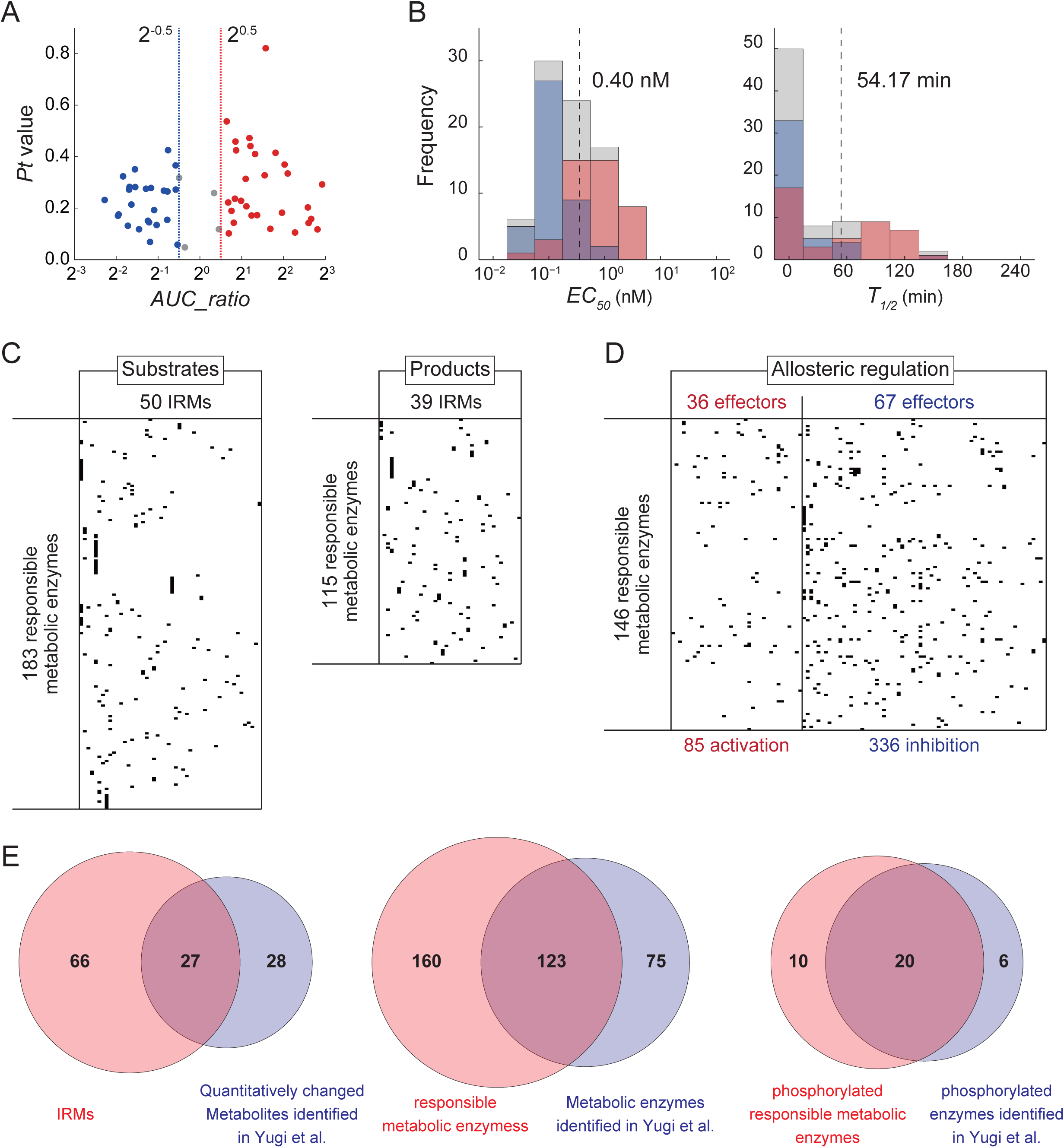
Identification of IRMs, responsible metabolic enzymes, and allosteric regulators, Related to Figure 4. (A) Distribution of *Pt* values and *AUC_ratios* calculated for the IRMs. Red and blue dots indicate increased and decreased IRMs, respectively. Gray dots indicate IRMs not included in neither increased nor decreased IRMs. (B) Distribution of *EC*_*50*_ and *T*_*1/2*_ values calculated for increased IRMs (red) and decreased IRMs (blue). Gray bars indicate sum of frequency of increased and decreased IRMs. (C) Substrates and products of the responsible metabolic enzymes. A dot indicates that the IRM (x-axis) is substrate or a product for a metabolic enzyme (y-axis). See TableS7. (D) Allosteric regulation of the responsible metabolic enzymes by the IRMs that function as positive (activation) or negative (inhibition) allosteric effectors. A dot indicates that the IRM (x-axis) is allosteric effector for a metabolic enzyme (y-axis). Information for allosteric regulation was obtained from the BRENDA database (see STAR Methods). (E) Comparison of IRMs, responsible metabolic enzymes, and phosphorylated responsible metabolic enzymes with the previous study (Yugi et al., 2014). Not all of the IRGs, IRMs, and responsible metabolic enzymes, in this study were overlapped with our previous study. The difference may relate to the differences in the insulin stimulation time and in how IRMs were defined. See Table S8.

**Figure S4.**
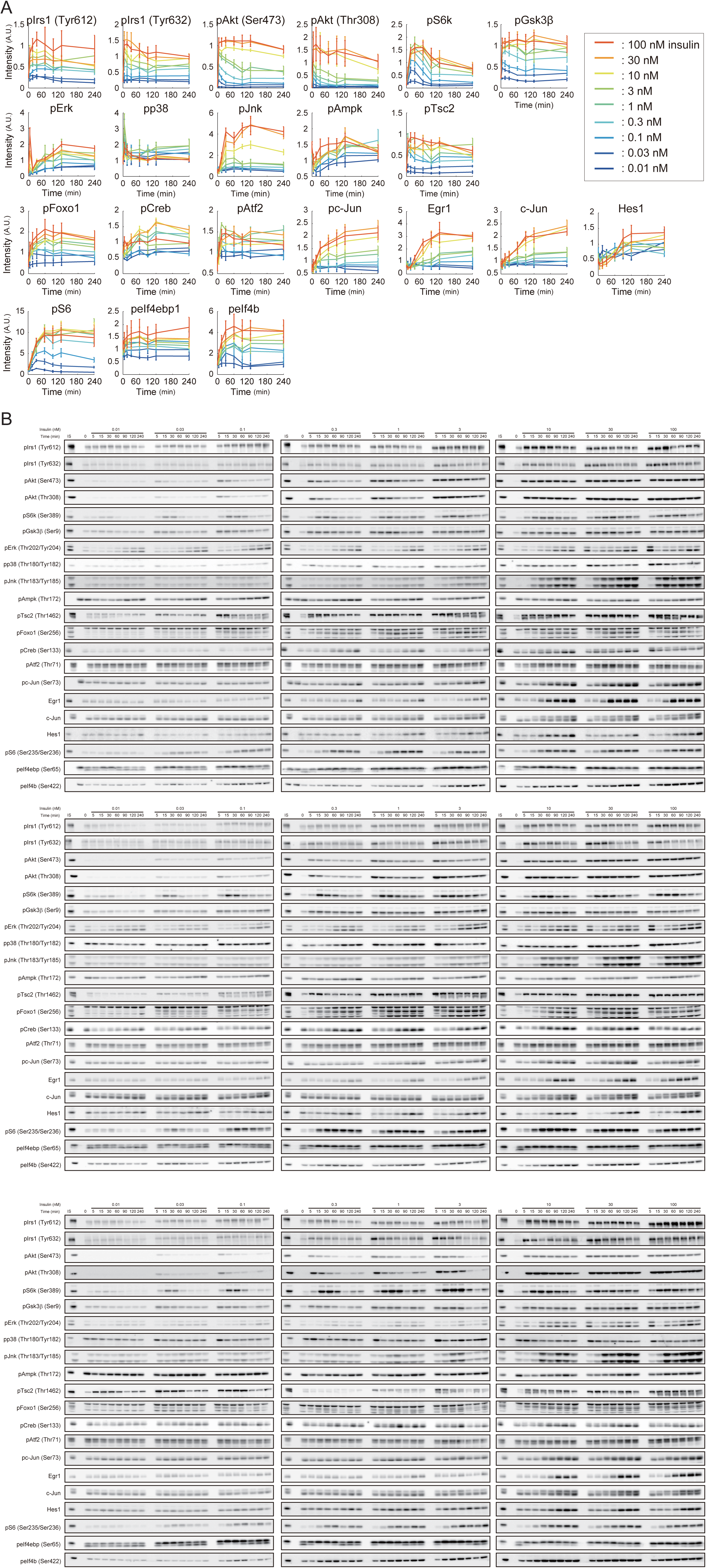
Time courses of the IRPs monitored by western blotting in response to insulin stimulation, Related to Figure 6. (A) Time courses of the indicated molecules by the indicated doses of insulin were plotted from data obtained by Western blotting. The means and SEMs of three independent experiments are shown. Lowercase ‘p’ preceding the name of a protein indicates the detection of the phosphorylated form of the protein. Numbers and letters in parentheses represent the phosphorylated amino acid residue. Numbering according to human. (B) All images of Western blotting were shown. IS indicates internal standard.

## STAR METHODS

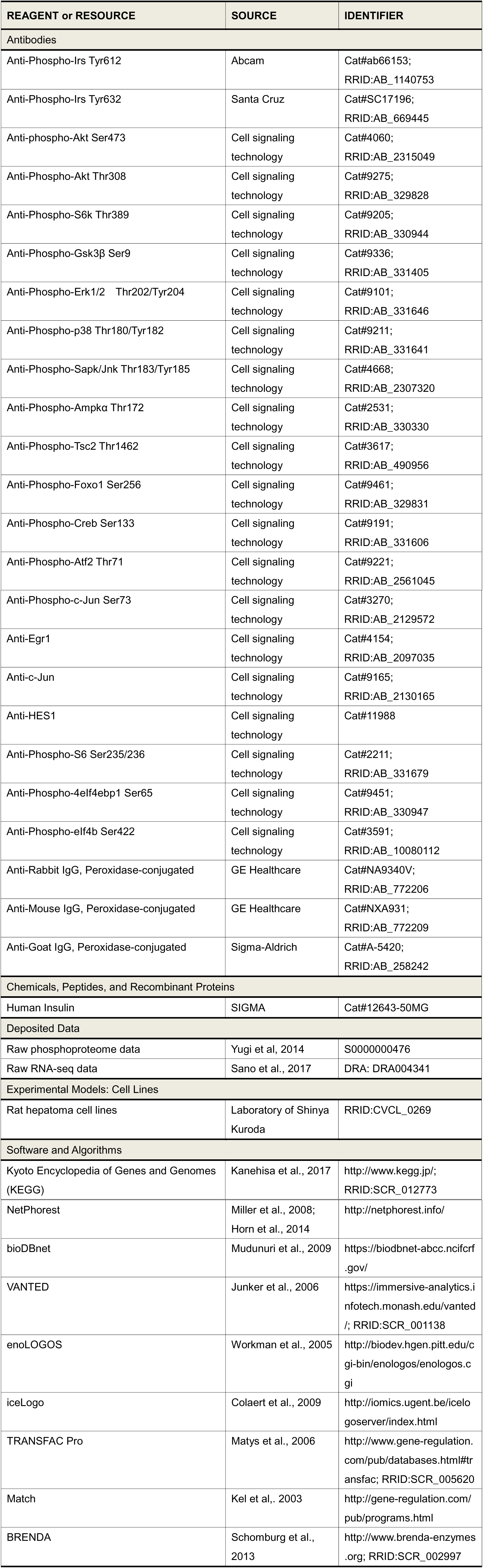
KEY RESOURCES TABLE

## CONTACT FOR REAGENT AND RESOURCE SHARING

Further information and requests for resources and reagents should be directed to and will be fulfilled by the Lead Contact, Shinya Kuroda.

## EXPERIMENTAL MODEL AND SUBJECT DETAILS

### Phosphoproteomic and Transcriptomic Data Sets

In this study, we used published datasets of the quantitative phosphoproteome (JPOST: S0000000476) (Yugi et al., 2014) and RNA-sequence (RNA-seq) (DDBJ: DRA004341) (Sano et al., 2016) of a time series of insulin stimulation of FAO cells (RRID:CVCL_0269, male). In the phosphoproteome analysis, FAO cells were stimulated with 1 nM insulin for 0, 2, 5, 10, 30, 45, and 60 min. Because the phosphoproteomic data consists of two different time series from two separate experiments (0, 5, 10, and 45 min and 2, 10, 30, and 60 min), some of the phosphopeptides were identified and quantified in data from only one of the time series. Therefore, a fold change of phosphorylation intensity was calculated as a ratio of the phosphorylation intensity at each time point to the phosphorylation intensity at t = 0 or 2 min. A phosphopeptide with a phosphorylation intensity greater than a 1.5-fold increase or less than a 0.67-fold decrease at more than one time point was defined as a quantitatively changed phosphopeptide. In the RNA-seq analysis, FAO cells were stimulated with 0.01, 1, and 100 nM insulin for 0, 15, 30, 60, 90, 120, and 240 min. The fragments per kilobase of transcript per million mapped reads (FPKM) values of each gene were calculated using Cufflinks (Trapnell et al., 2009, 2012), and the differentially expressed transcripts were identified using Cuffdiff (Trapnell et al., 2009, 2012). The genes encoding the differentially expressed transcripts were defined as IRGs.

### Sample Preparation for Metabolomic Analysis and Western Blotting

#### Sample preparation

Rat hepatoma FAO cells were seeded at a density of 3 × 10^6^ cells per dish on 6-cm dishes (Corning) or 1.3 × 10^6^ cells per well on six-well plates (Iwaki) and cultured in RPMI 1640 supplemented with 10% (v/v) fetal bovine serum at 37°C under 5% CO_2_ for 2 days before deprivation of serum (starvation). The cells were washed twice with phosphate-buffered saline (PBS) and starved for 16 hours in serum-free medium including 0.01 nM insulin (Sigma-Aldrich) and 10 nM dexamethasone (Wako), which increases the expression of gluconeogenesis genes such as *G6pase* and *Pck1* (Lange et al., 1994). We continuously added 0.01 nM insulin before the stimulation, and 0.01 nM insulin was present throughout the experiments unless otherwise specified to mimic the in vivo basal secretion during fasting (Polonsky et al., 1988). The medium was changed at 4 and 2 hours before the stimulation. Cells were stimulated with the indicated doses of insulin.

#### Metabolomic Analysis

For metabolomic analyses, the cells were washed at the indicated times after insulin stimulation with 4 mL ice-cold 5% mannitol twice and metabolites were extracted with 1 mL of ice-cold methanol that included the reference compounds [25 μM L-methionine sulfone (Wako), 25 μM 2-Morpholinoethanesulfonic acid, monohydrate (Dojindo), and 25 μM D-Camphor-10-sulfonic acid (Wako)] for normalization of peak intensities of mass spectrometry among samples. The resulting supernatant (400 μL) was sequentially mixed with 200 μL of water and 400 μL of chloroform and then centrifuged at 12,000 × *g* for 15 min at 4°C. The separated aqueous layer was filtered through a 5 kDa cutoff filter (Millipore) to remove proteins. The filtrate (320 μL) was lyophilized and dissolved in 50 μL water including reference compounds [200 μM each of trimesate (Wako) and 3-aminopyrrolidine (Sigma-Aldrich)] for migration time and then injected into the capillary electrophoresis time-of-flight mass spectrometry (CE-TOFMS) system (Agilent Technologies) (Ishii et al., 2007; Soga et al., 2006, 2009).

#### Western blotting

For Western blotting, the cells were washed with ice-cold PBS and proteins were extracted with 50 mM Tris-Cl pH 8.8 including 1% SDS at the indicated times after insulin stimulation. The lysates were sonicated and centrifuged at 12,000 × *g* at 4 °C for 15 min to remove debris. Total protein concentration of the resulting supernatants was determined with the bicinchoninic acid assay (Thermo Fisher Scientific) and adjusted to 0.75 mg/mL.

Equal amount of total protein was loaded for SDS-PAGE followed by Western blotting with the indicated antibodies. Band intensities were measured by using TotalLab Quant software (Nonlinear Inc.). Lysate mixture of FAO cells stimulated with or without 100 nM insulin for 5 min was used as an internal standard to normalize the band intensities for each membrane.

## METHOD DETAILS

### Step I: Construction of the Cellular Functions Regulatory Network

#### Step I-i: Construction of the signaling layer

We selected pathways from the KEGG database that included the character string of “signaling pathway” in their names for signaling pathways. We integrated the signaling pathways in which the IRpPs were significantly over-represented into a single signaling pathway, and then removed the proteins that did not have detectable transcripts in our previous study (Sano et al., 2016) or that are not located in downstream of InsR in the KEGG database. We defined this integrated signaling pathway as the signaling layer (Figure 2A). The signaling layer in *.gml format is also downloadable from our website (http://kurodalab.bs.s.u-tokyo.ac.jp/info/Kawata/). We recommend VANTED (https://immersive-analytics.infotech.monash.edu/vanted/) (Junker et al., 2006) to open the *.gml file.

#### Step I-ii: Construction of the cellular functional layer

We selected rat pathways from KEGG database for cellular functions with the following exceptions: (i) the signaling pathways (43 pathways); (ii) the global pathways (rno01XXX) including *Metabolic pathways* (rno01100) (9 pathways); (iii) the disease-related pathways (rno05XXX) (63 pathways); (iv) the pathways that work in specific tissues other than liver or are not regulated by insulin signaling in the liver (52 pathways); and (v) the pathways that include the character string of “diabetes”, “NAFLD”, or “Insulin resistance” in their names (8 pathways). The global pathways and disease-related pathways were excluded from cellular functional pathways because of redundancy with other cellular functional pathways as subsets. The cellular functional pathways in which the quantitatively changed phosphopeptides were significantly over-represented were defined as the cellular functional layer.

#### Step I-iii: Estimation of responsible protein kinases

We estimated kinase-substrate relationships for amino acid sequences of the proteins that had the quantitatively changed phosphopeptides using a stand-alone version of NetPhorest (http://netphorest.info/download/netphorest_human.tsv.xz) (Horn et al., 2014; Miller et al., 2008). The inputs for NetPhorest are rat protein sequences that are associated with international protein index (IPI) (Kersey et al., 2004) in FASTA format (ftp://ftp.ebi.ac.uk/pub/databases/IPI/current/ipi.RAT.fasta.gz). The outputs for NetPhorest are posterior probabilities of an amino acid residue being recognized by a protein kinase classifier (class of kinases with similar substrate recognition motifs). Among the candidate classifiers, we selected the classifier that had the largest posterior probability value as the kinase classifier related to the amino acid sequence. A predicted kinase-substrate relationship is represented as an edge between a kinase classifier as one node and a phosphorylation site of a phosphopeptide in an IRpP as the other node. Furthermore, we extracted individual kinases within in each class from NetPhorest (view-source:http://netphorest.info/download.shtml) and defined these kinases as responsible protein kinases.

### Step II: Construction of the Transcriptional Regulatory Network

#### Step II-i: Definition of up-regulated and down-regulated IRGs

The fold changes of FPKMs against those at 0 min were calculated for each IRG. The *log*_2_ of the fold changes were calculated to make the range of up-regulation and down-regulation comparable. The logarithms of fold changes were normalized between 0 and 1 to exclude the influence of constitutive expression. We defined the *Pt* value as an index of expression variation by taking the sum of the absolute values of the differences in the slopes at specific time points and at earlier or later time points, in response to 0.01 nM and 100 nM insulin stimulation (Figure S2A). A smaller *Pt* value indicates that the time series of gene expression has less variability. We defined *AUC_ratio* as an index of response by taking the ratio of AUC in response to 100 nM and that in response to 0.01 nM insulin (Figure S2A). The larger the absolute value of the *AUC_ratio* indicates that the response to insulin is larger. Here, genes with a *Pt* value larger than 0.2 were excluded from IRGs because of low quality of quantification. Among the IRGs with *Pt* values that were less than 0.2, those with an *AUC_ratio* of more than 2^0.5^ were defined as up-regulated IRGs, and those with an *AUC_ratio* of less than 2^-0.5^ were defined as down-regulated IRGs.

#### Step II-ii: Prediction of TFs regulating each class of IRGs

We predicted the TFs that regulate the expression of the classified IRGs by TF binding motif prediction and motif enrichment analysis. The flanking regions around the major transcription start site of each IRG were extracted from Rnor_5.0 (Ensembl, release 73) using Ensembl BioMart (Kinsella et al., 2011). We considered the genomic regions from -300 bp to +100 bp of the consensus transcription start sites as the flanking regions, according to the FANTOM5 time course analysis (Arner et al., 2015). We predicted the TF binding motifs that can bind to each flanking region using a TF database, TRANSFAC Pro (Matys et al., 2006), and Match, a TF binding motifs prediction tool. We used extended *vertebrate_non_redundant_min_SUM.prf*, one of the parameter sets prepared in TRANSFAC Pro for the threshold of similarity score calculated by Match. Because some of the TFs known to be regulated by insulin, including Foxo1, are not included in this parameter set, we extracted from *vertebrate_non_redundant.prf* the TF binding motifs that were not included in *vertebrate_non_redundant_min_SUM.prf* but were present in TFs included in KEGG *insulin signaling pathway* (rno4910), and we appended these TF binding motifs and their parameters to *vertebrate_non_redundant_min_SUM.prf*. The binding sites within each flanking region were predicted using Match with the extended *vertebrate_non_redundant_min_SUM.prf*. Enrichment of binding sites of TF binding motifs were determined using motif enrichment analysis in reference to CIDER methods (Mina et al., 2015), and the TFs related to significantly enriched TF binding motifs were identified as the TFs regulating IRGs in each class. An interactive network for the IRGs and the TFs in *.gml format is downloadable from our website (http://kurodalab.bs.s.u-tokyo.ac.jp/info/Kawata/). We recommend VANTED (https://immersive-analytics.infotech.monash.edu/vanted/) (Junker et al., 2006) to open the *.gml file.

#### Step II-iii: Identification of regulators of TFs

Using the accession numbers from TRANSFAC, we associated the significantly enriched TF binding motifs with TFs using the correspondence obtained from *matrix.dat* in TRANSFAC Pro. The accession numbers of TFs provided in TRANSFAC Pro are associated with the gene IDs for DATF, EMBL, FLYBASE, MIRBASE, PATHODB, PDB, SMARTDB, SWISSPROT, TRANSCOMPEL, or TRANSPATH. To identify regulators of the TFs, the gene IDs of EMBL, PDB, or SWISSPROT that were associated with the accession numbers of human, mouse, and rat TF were converted to KEGG gene IDs using bioDBnet (https://biodbnet-abcc.ncifcrf.gov/) (Mudunuri et al., 2009). We manually determined the upstream molecules of the TFs from the pathway information of KEGG, and except for those in the diseases related pathways (rno05XXX), these were defined as regulators.

### Step III: Construction of the Metabolism Regulatory Network

#### Step III-i: Identification of IRMs

We identified IRMs by comparing three factors: temporal changes of metabolites against the value at 0 min, changes in response to 0.01 and 100 nM insulin stimulation at each time point, and the data acquired on different days (*n*=3). Response of each metabolite to insulin concentrations was determined by three-way ANOVA with FDR using Storey’s procedure (Storey et al., 2004), and the metabolites showing a significant change (*FDR* < 0.1) were defined as IRMs.

#### Step III-ii: Definition of increased and decreased IRMs

Increased and decreased IRMs were defined using the same procedure as we used to identify the up-regulated and down-regulated IRGs. For each IRM, the fold change in the abundance of metabolites at each time point relative to the mean abundance at 0 min was calculated. We calculated *log*_2_ values of the fold changes so that the ranges of increased and decreased IRMs become comparable. The logarithmic values of fold change were normalized between 0 and 1, and the *AUC_ratio* was determined as the ratio of AUC with 100 nM insulin to AUC with 0.01 nM stimulation. The metabolites with an *AUC_ratio* of more than 2^0.5^ were defined as increased, and those with the *AUC_ratio* of less than 2^-0.5^ were defined as decreased IRMs.

#### Step III-iii: Identification of responsible metabolic enzymes

We searched the data of enzymatic reactions downloaded from the KEGG database. The IRMs, except for hub metabolites such as ATP and NADP^+^, were identified as targets of responsible metabolic enzymes. The hub metabolites were determined according to previous studies of network topology (Alves et al., 2002). Finally, we identified the enzymes with substrates or products that include at least one IRM, except the hub metabolites, and defined those as responsible metabolic enzymes.

#### Step III-iv: Identification of allosteric regulation

We identified allosteric regulation for each responsible metabolic enzyme using the same procedures used in the previous study (Yugi et al., 2014). We obtained the entries for the responsible metabolic enzymes from the BRENDA database (http://www.brenda-enzymes.org) (Schomburg et al., 2013) and extracted their allosteric effector (activator and inhibitor) information, as reported for mammals (*Bos Taurus, Felis catus, Homo sapiens*, “Macaca”, “Mammalia”, “Monkey”, *Mus booduga, Mus musculus, Rattus norvegicus, Rattus rattus, Rattus* sp., *Sus scrofa*, “dolphin”, and “hamster”). Then, we associated the standard compound names of allosteric effectors used in BRENDA with metabolite names that were used in KEGG to obtain the KEGG compound ID related to each allosteric effector. With this information, we matched the allosteric effectors included in IRMs to the responsible metabolic enzymes.

## QUANTIFICATION AND STATISTICAL ANALYSIS

### Pathway Over-Representation Analysis for Phosphoproteomic Data

Over-representation analysis with phosphoproteomic data was performed to identify the pathways enriched in the IRpPs. Signaling and cellular functional pathways were used for the over-representation analysis. The IDs of phosphopeptides provided as IPI (Kersey et al., 2004) in the phosphoproteomic data were converted to KEGG gene ID using bioDBnet (Mudunuri et al., 2009) to correspond with the pathway information of KEGG database.

Over-representation of the IRpPs for each pathway was determined by Fisher’s exact test (Fisher, 1922) with FDR using Storey’s procedure (Storey et al., 2004). This analysis identified the pathways indicating significant over-representation (*FDR* < 0.1).

### Clustering the Cellular Functions Pathways and Metabolic Enzymes by the Occurrence Rates of Responsible Protein Kinases

We performed hierarchical clustering analysis on the cellular functional pathways included in the cellular functional layer and metabolic enzymes included in the KEGG database using the occurrence rate of the predicted responsible protein kinases. The occurrence rate of a specific kinase classifier (*i*) in a specific pathway (*j*) was calculated as the ratio of the number of quantitatively changed phosphopeptides having the kinase classifier *i* including the responsible proteins kinases to the total number of quantitatively changed phosphopeptides in the pathways *j*. The sum of occurrence rates of the kinase classifiers in pathway *j* is 1. We performed the hierarchical clustering of the cellular functions pathways using the Euclidean distance for calculation of the intracluster distances and using Ward’s method for calculation of the intercluster distances (Ward, 1963).

### Generating Motif Logos

Motif logos of quantitatively changed phosphopeptides included in each cellular functional pathway and the metabolic enzymes were generated using enoLOGOS (http://biodev.hgen.pitt.edu/enologos/) (Workman et al., 2005) with relative entropy as logo plot methods. Ranges of motifs were provided as from -5 to +5 residues from phosphorylated sites. Statistical tests of amino acid compositions at each position of the quantitatively changed phosphopeptides included in cellular functions and metabolic enzymes, or each cluster were performed using iceLogo (http://iomics.ugent.be/icelogoserver/index.html) (Colaert et al., 2009) with percentage difference as scoring system and *p* value cut-off of 0.05. The *Rattus norvegicus* amino acid compositions from Swiss-Prot and quantitatively changed phosphopeptides included in cellular functions and metabolic enzymes were used as reference compositions for tests of quantitatively changed phosphopeptides included in cellular functions and metabolic enzymes and for tests of those included in each cluster.

### Motif Enrichment Analysis

Motif enrichment analysis in reference to CIDER methods (Mina et al., 2015) was performed to determine enrichment of binding sites of TF binding motifs in flanking region of IRGs in each class. Binding sites of each TF binding motif were determined using flanking regions obtained from Ensembl BioMart (Kinsella et al., 2011), TRANSFAC Pro (Matys et al., 2006), and Match tool. The enrichment of TF binding motif binding sites in the flanking regions of IRGs in each class were determined by Fisher’s exact test (Fisher, 1922) with FDR using Storey’s procedure (Storey et al., 2004). The TFs related to significantly enriched TF binding motifs (*FDR* < 0.1) were identified as the TFs regulating IRGs in each class.

### Classification of Signaling Molecules, IRGs, and IRMs

#### Estimation of sensitivities and response time by calculation of *EC*_*50*_ and *T*_*1/2*_

*EC*_*50*_ was defined as the concentration of insulin that gives the 50% of the maximal AUC of time series of responses (Figure S2D). A smaller *EC*_*50*_ indicates a higher sensitivity to insulin concentration. To exclude the influence of the difference in time series, we used the AUC of the time courses in response to each concentration of insulin to calculate *EC*_*50*_. *T*_*1/2*_ was defined as the time when the response reached 50% of the peak amplitude (Figure S2D). A smaller *T*_*1/2*_ indicates a faster response. The *T*_*1/2*_ values for the IRGs, the IRMs, and IRPs were estimated from the time course in response to 100 nM insulin stimulation. The distribution of *EC*_*50*_ and *T*_*1/2*_ values for IRGs under various thresholds of Cuffdiff (*FDR* < 0.01, 0.03, 0.05, 0.07, and 0.10; default: *FDR* < 0.05) were compared, to confirm that the distributions of the *EC*_*50*_ and the *T*_*1/2*_ were stable. The distributions of *EC*_*50*_ and *T*_*1/2*_ values between the up-regulated and down-regulated IRGs, or between the increased and decreased IRMs, were compared using Wilcoxon rank sum test (Gibbons and Chakraborti, 2011; Hollander et al., 2015). The *p* values were adjusted for multiple testing with Bonferroni correction (Bonferroni, 1936).

#### Definition of classes of IRPs, IRGs, and IRMs

To characterize the IRPs, the IRGs, or the IRMs by sensitivities (*EC*_*50*_) and response time (*T*_*1/2*_) to insulin stimulation, we classified these molecules using the *EC*_*50*_ and the *T*_*1/2*_ values. The thresholds dividing fast or slow responses and high or low sensitivity were determined using Otsu’s method (Otsu, 1979). The thresholds of the *EC*_*50*_ and the *T*_*1/2*_ values for the IRPs were determined from Western blotting data. The thresholds of the *EC*_*50*_ and the *T*_*1/2*_ values for the IRGs and the IRMs were determined using transcriptomic and metabolomic data, respectively. Using the thresholds, we divided the distributions of *EC*_*50*_ values into high and low sensitivities, and those of *T*_*1/2*_ values into fast and slow responses. We classified the signaling factors, the TFs, the IRGs, and the IRMs into four classes according to the thresholds: Class 1, high sensitivity (*EC*_*50*_ < threshold) and fast response (*T*_*1/2*_ < threshold) and; Class 2, high sensitivity and slow response (*T*_*1/2*_ > threshold); Class 3, low sensitivity (*EC*_*50*_ > threshold) and fast response, and Class 4, low sensitivity and slow response.

### Three-way ANOVA

Three-way ANOVA was performed to identify IRMs by comparing three factors: temporal changes of metabolites against the value at 0 min, responses to 0.01 and 100 nM insulin stimulation at each time points, and the data acquired on different days (*n*=3). The fold change of abundance of metabolites relative to the mean abundance at 0 min was calculated for each metabolite. We calculated *log*_2_ values of the fold changes so that ranges of increased and decreased IRMs become comparable. We performed three-way ANOVA with insulin doses (0.01 and 100 nM), time points after insulin stimulation (0, 5, 15, 30, 60, 90, 120, and 240 min), and data sets using the logarithmic values of fold changes. The *p* values against insulin concentrations were calculated and the FDR for each metabolite was calculated by Storey’s procedures (Storey et al., 2004). The λ value to calculate FDR was set to 0.8 with reference to the distribution of *p* values. The metabolites showing significance (*FDR* < 0.1) were defined as IRMs.

## DATA AND SOFTWARE AVAILABILITY

The raw phosphoproteomic data generated in previous study (Yugi et al., 2014) and used in this study have been deposited in the JPOST under ID code S0000000476. The raw transcriptomic data generated in previous study (Sano et al., 2016) and used in this study have been deposited in the DDBJ under ID code DRA004341.

## SUPPLEMENTAL ITEMS

Table S1. Pathway over-representation analysis, Related Figure 2.

Table S2. Prediction of responsible protein kinases, Related Figure 2.

Table S3. Occurrence rates of kinase classifiers in each cellular functional pathway, Related Figure 2.

Table S4. Classification of insulin-responsive genes (IRGs), Related to Figure 3 and 6.

Table S5. Prediction of transcription factors (TFs) for each class of insulin-responsive genes (IRGs), Related to Figure 3.

Table S6. Time series of metabolome data in response to insulin stimulation, Related to Figure 4.

Table S7. Classification of insulin-responsive metabolites (IRMs), Related to Figure 4 and 6.

Table S8. Identification of responsible metabolic enzymes, Related to Figure 4.

Table S9. Identification of allosteric regulators, Related to Figure 4.

Table S10. Classification of insulin-responsive proteins (IRPs), Related to Figure 6.

## REFERENCES

Aiston, S., Hampson, L.J., Arden, C., Iynedjian, P.B., and Agius, L. (2006). The role of protein kinase B/Akt in insulin-induced inactivation of phosphorylase in rat hepatocytes. Diabetologia 49, 174–182.

Alves, R., Chaleil, R.A.G., and Sternberg, M.J.E. (2002). Evolution of enzymes in metabolism: a network perspective. J. Mol. Biol. 320, 751–770.

Arner, E., Daub, C.O., Vitting-Seerup, K., Andersson, R., Lilje, B., Drabløs, F., Lennartsson, A., Rönnerblad, M., Hrydziuszko, O., Vitezic, M., et al. (2015). Transcribed enhancers lead waves of coordinated transcription in transitioning mammalian cells. Science 347, 1010–1014.

Asnaghi, L., Bruno, P., Priulla, M., and Nicolin, A. (2004). mTOR: a protein kinase switching between life and death. Pharmacol. Res. 50, 545–549.

Au, W.-S., Kung, H., and Lin, M.C. (2003). Regulation of microsomal triglyceride transfer protein gene by insulin in HepG2 cells: roles of MAPKerk and MAPKp38. Diabetes 52, 1073–1080.

Barthel, A., Schmoll, D., and Unterman, T.G. (2005). FoxO proteins in insulin action and metabolism. Trends Endocrinol. Metab. 16, 183–189.

Behar, M., and Hoffmann, A. (2010). Understanding the temporal codes of intra-cellular signals. Curr. Opin. Genet. Dev. 20, 684–693.

Biggs, W.H., Meisenhelder, J., Hunter, T., Cavenee, W.K., and Arden, K.C. (1999). Protein kinase B/Akt-mediated phosphorylation promotes nuclear exclusion of the winged helix transcription factor FKHR1. Proc. Natl. Acad. Sci. U. S. A. 96, 7421–7426.

Bonferroni (1936). Teoria statistica delle classi e calcolo delle probabilità. Pubbl. Del R Ist. Super. Di Sci. Econ. E Commer. Di Firenze 8.

Brabant, G., Prank, K., and Schofl, C. (1992). Pulsatile patterns in hormone secretion. Trends Endocrinol. Metab. 3, 183–190.

Brunet, A., Bonni, A., Zigmond, M.J., Lin, M.Z., Juo, P., Hu, L.S., Anderson, M.J., Arden, K.C., Blenis, J., and Greenberg, M.E. (1999). Akt promotes cell survival by phosphorylating and inhibiting a Forkhead transcription factor. Cell 96, 857–868.

Buescher, J.M., Liebermeister, W., Jules, M., Uhr, M., Muntel, J., Botella, E., Hessling, B., Kleijn, R.J., Le Chat, L., Lecointe, F., et al. (2012). Global Network Reorganization During Dynamic Adaptations of Bacillus subtilis Metabolism. Science (80-. ). 335, 1099–1103.

Chiappino-Pepe, A., Pandey, V., Ataman, M., and Hatzimanikatis, V. (2017). Integration of metabolic, regulatory and signaling networks towards analysis of perturbation and dynamic responses. Curr. Opin. Syst. Biol. 2, 59–66.

Colaert, N., Helsens, K., Martens, L., Vandekerckhove, J., and Gevaert, K. (2009). Improved visualization of protein consensus sequences by iceLogo. Nat. Methods 6, 786–787.

Deak, M., Clifton, A.D., Lucocq, L.M., and Alessi, D.R. (1998). Mitogen‐ and stress-activated protein kinase-1 (MSK1) is directly activated by MAPK and SAPK2/p38, and may mediate activation of CREB. EMBO J. 17, 4426–4441.

Dupont, J., Khan, J., Qu, B.-H., Metzler, P., Helman, L., and LeRoith, D. (2001). Insulin and IGF-1 Induce Different Patterns of Gene Expression in Mouse Fibroblast NIH-3T3 Cells: Identification by cDNA Microarray Analysis. Endocrinology 142, 4969–4975.

Durr, I.F., and Rudney, H. (1960). The reduction of beta-hydroxy-beta-methyl-glutaryl coenzyme A to mevalonic acid. J. Biol. Chem. 235, 2572–2578.

Everman, S., Meyer, C., Tran, L., Hoffman, N., Carroll, C.C., Dedmon, W.L., and Katsanos, C.S. (2016). Insulin does not stimulate muscle protein synthesis during increased plasma branched-chain amino acids alone but still decreases whole body proteolysis in humans. Am. J. Physiol. Endocrinol. Metab. 311, E671–E677.

Fisher, R.A. (1922). On the Interpretation of χ 2 from Contingency Tables, and the Calculation of P. J. R. Stat. Soc. 85, 87.

Friedman, A.A., Tucker, G., Singh, R., Yan, D., Vinayagam, A., Hu, Y., Binari, R., Hong, P., Sun, X., Porto, M., et al. (2011). Proteomic and Functional Genomic Landscape of Receptor Tyrosine Kinase and Ras to Extracellular Signal-Regulated Kinase Signaling. Sci. Signal. 4, rs10–rs10.

Fukunaga, K., Noguchi, T., Takeda, H., Matozaki, T., Hayashi, Y., Itoh, H., and Kasuga, M. (2000). Requirement for protein-tyrosine phosphatase SHP-2 in insulin-induced activation of c-Jun NH(2)-terminal kinase. J. Biol. Chem. 275, 5208–5213.

Geiger, R., Rieckmann, J.C., Wolf, T., Basso, C., Feng, Y., Fuhrer, T., Kogadeeva, M., Picotti, P., Meissner, F., Mann, M., et al. (2016). L-Arginine Modulates T Cell Metabolism and Enhances Survival and Anti-tumor Activity. Cell 167, 829–842.e13.

Gerosa, L., Haverkorn van Rijsewijk, B.R.B., Christodoulou, D., Kochanowski, K., Schmidt, T.S.B., Noor, E., and Sauer, U. (2015). Pseudo-transition Analysis Identifies the Key Regulators of Dynamic Metabolic Adaptations from Steady-State Data. Cell Syst. 1, 270–282.

Gibbons, J.D., and Chakraborti, S. (2011). Nonparametric statistical inference (Chapman & Hall/Taylor & Francis).

Gonçalves, E., Raguz Nakic, Z., Zampieri, M., Wagih, O., Ochoa, D., Sauer, U., Beltrao, P., and Saez-Rodriguez, J. (2017). Systematic Analysis of Transcriptional and Post-transcriptional Regulation of Metabolism in Yeast. PLOS Comput. Biol. 13, e1005297.

Hackett, S.R., Zanotelli, V.R.T., Xu, W., Goya, J., Park, J.O., Perlman, D.H., Gibney, P.A., Botstein, D., Storey, J.D., and Rabinowitz, J.D. (2016). Systems-level analysis of mechanisms regulating yeast metabolic flux. Science (80-. ). 354, aaf2786–aaf2786.

Hartmann, B., Castelo, R., Blanchette, M., Boue, S., Rio, D.C., and Valcárcel, J. (2009). Global analysis of alternative splicing regulation by insulin and wingless signaling in Drosophila cells. Genome Biol. 10, R11.

Hatzimanikatis, V., and Saez-Rodriguez, J. (2015). Integrative approaches for signalling and metabolic networks. Integr. Biol. (Camb). 7, 844–845.

Hectors, T.L.M., Vanparys, C., Pereira-Fernandes, A., Knapen, D., and Blust, R. (2012). Mechanistic evaluation of the insulin response in H4IIE hepatoma cells: New endpoints for toxicity testing? Toxicol. Lett. 212, 180–189.

Herzig, S., Hedrick, S., Morantte, I., Koo, S.-H., Galimi, F., and Montminy, M. (2003). CREB controls hepatic lipid metabolism through nuclear hormone receptor PPAR-γ. Nature 426, 190–193.

Hollander, M., A. Wolfe, D., and Chicken, E. (2015). Nonparametric Statistical Methods (Hoboken, NJ, USA: John Wiley & Sons, Inc.).

Horn, H., Schoof, E.M., Kim, J., Robin, X., Miller, M.L., Diella, F., Palma, A., Cesareni, G., Jensen, L.J., and Linding, R. (2014). KinomeXplorer: an integrated platform for kinome biology studies. Nat. Methods 11, 603–604.

Humphrey, S., Yang, G., Yang, P., Fazakerley, D., St?ckli, J., Yang, J., and James, D. (2013). Dynamic Adipocyte Phosphoproteome Reveals that Akt Directly Regulates mTORC2. Cell Metab. 17, 1009–1020.

Humphrey, S.J., Azimifar, S.B., and Mann, M. (2015). High-throughput phosphoproteomics reveals in vivo insulin signaling dynamics. Nat. Biotechnol. 33, 990–995.

Hyduke, D.R., Lewis, N.E., and Palsson, B.Ø. (2013). Analysis of omics data with genome-scale models of metabolism. Mol. Biosyst. 9, 167–174.

Ishii, N., Nakahigashi, K., Baba, T., Robert, M., Soga, T., Kanai, A., Hirasawa, T., Naba, M., Hirai, K., Hoque, A., et al. (2007). Multiple high-throughput analyses monitor the response of E. coli to perturbations. Science 316, 593–597.

Jastrzebski, K., Hannan, K.M., Tchoubrieva, E.B., Hannan, R.D., and Pearson, R.B. (2007). Coordinate regulation of ribosome biogenesis and function by the ribosomal protein S6 kinase, a key mediator of mTOR function. Growth Factors 25, 209–226.

Joyce, A.R., and Palsson, B.Ø. (2006). The model organism as a system: integrating “omics” data sets. Nat. Rev. Mol. Cell Biol. 7, 198–210.

Junker, B.H., Klukas, C., and Schreiber, F. (2006). VANTED: a system for advanced data analysis and visualization in the context of biological networks. BMC Bioinformatics 7, 109.

Kanehisa, M., Goto, S., Sato, Y., Furumichi, M., and Tanabe, M. (2012). KEGG for integration and interpretation of large-scale molecular data sets. Nucleic Acids Res. 40, D109–14.

Kanehisa, M., Furumichi, M., Tanabe, M., Sato, Y., and Morishima, K. (2017). KEGG: new perspectives on genomes, pathways, diseases and drugs. Nucleic Acids Res. 45, D353–D361.

Karoor, V., Wang, L., Wang, H.Y., and Malbon, C.C. (1998). Insulin stimulates sequestration of beta-adrenergic receptors and enhanced association of beta-adrenergic receptors with Grb2 via tyrosine 350. J. Biol. Chem. 273, 33035–33041.

Kel, A.E., Gössling, E., Reuter, I., Cheremushkin, E., Kel-Margoulis, O. V, and Wingender, E. (2003). MATCH: A tool for searching transcription factor binding sites in DNA sequences. Nucleic Acids Res. 31, 3576–3579.

Kersey, P.J., Duarte, J., Williams, A., Karavidopoulou, Y., Birney, E., and Apweiler, R. (2004). The International Protein Index: An integrated database for proteomics experiments. Proteomics 4, 1985–1988.

Kim, H.S., and Lee, N.K. (2014). Gene expression profiling in osteoclast precursors by insulin using microarray analysis. Mol. Cells 37, 827–832.

Kinsella, R.J., Kahari, A., Haider, S., Zamora, J., Proctor, G., Spudich, G., Almeida-King, J., Staines, D., Derwent, P., Kerhornou, A., et al. (2011). Ensembl BioMarts: a hub for data retrieval across taxonomic space. Database 2011, bar030–bar030.

Kotzka, J., Lehr, S., Roth, G., Avci, H., Knebel, B., and Muller-Wieland, D. (2004). Insulin-activated Erk-mitogen-activated protein kinases phosphorylate sterol regulatory element-binding Protein-2 at serine residues 432 and 455 in vivo. J. Biol. Chem. 279, 22404–22411.

Krüger, M., Kratchmarova, I., Blagoev, B., Tseng, Y.-H., Kahn, C.R., and Mann, M. (2008). Dissection of the insulin signaling pathway via quantitative phosphoproteomics. Proc. Natl. Acad. Sci. U. S. A. 105, 2451–2456.

Kubota, H., Noguchi, R., Toyoshima, Y., Ozaki, Y., Uda, S., Watanabe, K., Ogawa, W., and Kuroda, S. (2012). Temporal Coding of Insulin Action through Multiplexing of the AKT Pathway. Mol. Cell 46, 820–832.

Lange, A.J., Argaud, D., el-Maghrabi, M.R., Pan, W., Maitra, S.R., and Pilkis, S.J. (1994). Isolation of a cDNA for the catalytic subunit of rat liver glucose-6-phosphatase: regulation of gene expression in FAO hepatoma cells by insulin, dexamethasone and cAMP. Biochem. Biophys. Res. Commun. 201, 302–309.

Lee, Y.H., Giraud, J., Davis, R.J., and White, M.F. (2003). c-Jun N-terminal kinase (JNK) mediates feedback inhibition of the insulin signaling cascade. J. Biol. Chem. 278, 2896–2902.

Lemke, U., Krones-Herzig, A., Diaz, M.B., Narvekar, P., Ziegler, A., Vegiopoulos, A., Cato, A.C.B., Bohl, S., Klingmüller, U., Screaton, R.A., et al. (2008). The Glucocorticoid Receptor Controls Hepatic Dyslipidemia through Hes1. Cell Metab. 8, 212–223.

Lindgren, V., Luskey, K.L., Russell, D.W., and Francke, U. (1985). Human genes involved in cholesterol metabolism: chromosomal mapping of the loci for the low density lipoprotein receptor and 3-hydroxy-3-methylglutaryl-coenzyme A reductase with cDNA probes. Proc. Natl. Acad. Sci. U. S. A. 82, 8567–8571.

Lindsay, J.R., McKillop, A.M., Mooney, M.H., Flatt, P.R., Bell, P.M., and O’Harte, F.P.M. (2003). Meal-induced 24-hour profile of circulating glycated insulin in type 2 diabetic subjects measured by a novel radioimmunoassay. Metabolism 52, 631–635.

Lizcano, J.M., and Alessi, D.R. (2002). The insulin signalling pathway. Curr. Biol. 12, R236–8.

Lu, M., Wan, M., Leavens, K.F., Chu, Q., Monks, B.R., Fernandez, S., Ahima, R.S., Ueki, K., Kahn, C.R., and Birnbaum, M.J. (2012). Insulin regulates liver metabolism in vivo in the absence of hepatic Akt and Foxo1. Nat. Med. 18, 388–395.

Matsuzaki, H., Daitoku, H., Hatta, M., Tanaka, K., and Fukamizu, A. (2003). Insulin-induced phosphorylation of FKHR (Foxo1) targets to proteasomal degradation. Proc. Natl. Acad. Sci. U. S. A. 100, 11285–11290.

Matys, V., Kel-Margoulis, O. V, Fricke, E., Liebich, I., Land, S., Barre-Dirrie, A., Reuter, I., Chekmenev, D., Krull, M., Hornischer, K., et al. (2006). TRANSFAC(R) and its module TRANSCompel(R): transcriptional gene regulation in eukaryotes. Nucleic Acids Res. 34, D108–D110.

Miller, M.L., Jensen, L.J., Diella, F., Jorgensen, C., Tinti, M., Li, L., Hsiung, M., Parker, S.A., Bordeaux, J., Sicheritz-Ponten, T., et al. (2008). Linear Motif Atlas for Phosphorylation-Dependent Signaling. Sci. Signal. 1, ra2–ra2.

Mina, M., Magi, S., Jurman, G., Itoh, M., Kawaji, H., Lassmann, T., Arner, E., Forrest, A.R.R., Carninci, P., Hayashizaki, Y., et al. (2015). Promoter-level expression clustering identifies time development of transcriptional regulatory cascades initiated by ErbB receptors in breast cancer cells. Sci. Rep. 5, 11999.

Monetti, M., Nagaraj, N., Sharma, K., and Mann, M. (2011). Large-scale phosphosite quantification in tissues by a spike-in SILAC method. Nat. Methods 8, 655–658.

Mounier, C., and Posner, B.I. (2006). Transcriptional regulation by insulin: from the receptor to the geneThis paper is one of a selection of papers published in this Special issue, entitled Second Messengers and Phosphoproteins—12th International Conference. Can. J. Physiol. Pharmacol. 84, 713–724.

Mudunuri, U., Che, A., Yi, M., and Stephens, R.M. (2009). bioDBnet: the biological database network. Bioinformatics 25, 555–556.

Murphy, L.O., MacKeigan, J.P., and Blenis, J. (2004). A network of immediate early gene products propagates subtle differences in mitogen-activated protein kinase signal amplitude and duration. Mol. Cell. Biol. 24, 144–153.

Nakayama, K., Satoh, T., Igari, A., Kageyama, R., and Nishida, E. (2008). FGF induces oscillations of Hes1 expression and Ras/ERK activation. Curr. Biol. 18, R332–4.

Noguchi, R., Kubota, H., Yugi, K., Toyoshima, Y., Komori, Y., Soga, T., and Kuroda, S. (2013). The selective control of glycolysis, gluconeogenesis and glycogenesis by temporal insulin patterns. Mol. Syst. Biol. 9, 664.

O’Meara, N.M., Sturis, J., Blackman, J.D., Roland, D.C., Van Cauter, E., and Polonsky, K.S. (1993). Analytical problems in detecting rapid insulin secretory pulses in normal humans. Am. J. Physiol. 264, E231–8.

O’Rahilly, S., Turner, R.C., and Matthews, D.R. (1988). Impaired Pulsatile Secretion of Insulin in Relatives of Patients with Non-Insulin-Dependent Diabetes. N. Engl. J. Med. 318, 1225–1230.

Oliveira, A.P., Ludwig, C., Picotti, P., Kogadeeva, M., Aebersold, R., and Sauer, U. (2012). Regulation of yeast central metabolism by enzyme phosphorylation. Mol. Syst. Biol. 8, 623.

Otsu, N. (1979). A Threshold Selection Method from Gray-Level Histograms. IEEE Trans. Syst. Man. Cybern. 9, 62–66.

Palsson, B., and Zengler, K. (2010). The challenges of integrating multi-omic data sets. Nat. Chem. Biol. 6, 787–789.

Polonsky, K.S., Given, B.D., and Van Cauter, E. (1988). Twenty-four-hour profiles and pulsatile patterns of insulin secretion in normal and obese subjects. J. Clin. Invest. 81, 442–448.

Puigserver, P., Rhee, J., Donovan, J., Walkey, C.J., Yoon, J.C., Oriente, F., Kitamura, Y., Altomonte, J., Dong, H., Accili, D., et al. (2003). Insulin-regulated hepatic gluconeogenesis through FOXO1-PGC-1alpha interaction. Nature 423, 550–555.

Purvis, J.E., and Lahav, G. (2013). Encoding and Decoding Cellular Information through Signaling Dynamics. Cell 152, 945–956.

Reiss, K., Wang, J.Y., Romano, G., Tu, X., Peruzzi, F., and Baserga, R. (2001). Mechanisms of regulation of cell adhesion and motility by insulin receptor substrate-1 in prostate cancer cells. Oncogene 20, 490–500.

Rome, S., Clément, K., Rabasa-Lhoret, R., Loizon, E., Poitou, C., Barsh, G.S., Riou, J.-P., Laville, M., and Vidal, H. (2003). Microarray profiling of human skeletal muscle reveals that insulin regulates approximately 800 genes during a hyperinsulinemic clamp. J. Biol. Chem. 278, 18063–18068.

Rosner, M., Siegel, N., Valli, A., Fuchs, C., and Hengstschl?ger, M. (2010). mTOR phosphorylated at S2448 binds to raptor and rictor. Amino Acids 38, 223–228.

Saltiel, A.R., and Kahn, C.R. (2001). Insulin signalling and the regulation of glucose and lipid metabolism. Nature 414, 799–806.

Sano, T., Kawata, K., Ohno, S., Yugi, K., Kakuda, H., Kubota, H., Uda, S., Fujii, M., Kunida, K., Hoshino, D., et al. (2016). Selective control of up-regulated and down-regulated genes by temporal patterns and doses of insulin. Sci. Signal. 112, 1–12.

Schomburg, I., Chang, A., Placzek, S., Söhngen, C., Rother, M., Lang, M., Munaretto, C., Ulas, S., Stelzer, M., Grote, A., et al. (2013). BRENDA in 2013: integrated reactions, kinetic data, enzyme function data, improved disease classification: new options and contents in BRENDA. Nucleic Acids Res. 41, D764–72.

Shaul, Y.D., and Seger, R. (2007). The MEK/ERK cascade: From signaling specificity to diverse functions. Biochim. Biophys. Acta - Mol. Cell Res. 1773, 1213–1226.

Skolnik, E.Y., Batzer, A., Li, N., Lee, C.H., Lowenstein, E., Mohammadi, M., Margolis, B., and Schlessinger, J. (1993a). The function of GRB2 in linking the insulin receptor to Ras signaling pathways. Science 260, 1953–1955.

Skolnik, E.Y., Lee, C.H., Batzer, A., Vicentini, L.M., Zhou, M., Daly, R., Myers, M.J., Backer, J.M., Ullrich, A., and White, M.F. (1993b). The SH2/SH3 domain-containing protein GRB2 interacts with tyrosine-phosphorylated IRS1 and Shc: implications for insulin control of ras signalling. EMBO J. 12, 1929–1936.

Soga, T., Baran, R., Suematsu, M., Ueno, Y., Ikeda, S., Sakurakawa, T., Kakazu, Y., Ishikawa, T., Robert, M., Nishioka, T., et al. (2006). Differential metabolomics reveals ophthalmic acid as an oxidative stress biomarker indicating hepatic glutathione consumption. J. Biol. Chem. 281, 16768–16776.

Soga, T., Igarashi, K., Ito, C., Mizobuchi, K., Zimmermann, H.-P., and Tomita, M. (2009). Metabolomic Profiling of Anionic Metabolites by Capillary Electrophoresis Mass Spectrometry. Anal. Chem. 81, 6165–6174.

Srivastava, A.K., and Pandey, S.K. (1998). Potential mechanism(s) involved in the regulation of glycogen synthesis by insulin. Mol. Cell. Biochem. 182, 135–141.

Storey, J.D., Taylor, J.E., and Siegmund, D. (2004). Strong control, conservative point estimation and simultaneous conservative consistency of false discovery rates: a unified approach. J. R. Stat. Soc. Ser. B (Statistical Methodol. 66, 187–205.

Sukhatme, V.P., Cao, X.M., Chang, L.C., Tsai-Morris, C.H., Stamenkovich, D., Ferreira, P.C., Cohen, D.R., Edwards, S.A., Shows, T.B., and Curran, T. (1988). A zinc finger-encoding gene coregulated with c-fos during growth and differentiation, and after cellular depolarization. Cell 53, 37–43.

Thiel, G., and Cibelli, G. (2002). Regulation of life and death by the zinc finger transcription factor Egr-1. J. Cell. Physiol. 193, 287–292.

Trapnell, C., Pachter, L., and Salzberg, S.L. (2009). TopHat: discovering splice junctions with RNA-Seq. Bioinformatics 25, 1105–1111.

Trapnell, C., Roberts, A., Goff, L., Pertea, G., Kim, D., Kelley, D.R., Pimentel, H., Salzberg, S.L., Rinn, J.L., and Pachter, L. (2012). Differential gene and transcript expression analysis of RNA-seq experiments with TopHat and Cufflinks. Nat. Protoc. 7, 562–578.

Tsakiridis, T., Tong, P., Matthews, B., Tsiani, E., Bilan, P.J., Klip, A., and Downey, G.P. (1999). Role of the actin cytoskeleton in insulin action. Microsc. Res. Tech. 47, 79–92.

Versteyhe, S., Klaproth, B., Borup, R., Palsgaard, J., Jensen, M., Gray, S.G., and De Meyts, P. (2013). IGF-I, IGF-II, and Insulin Stimulate Different Gene Expression Responses through Binding to the IGF-I Receptor. Front. Endocrinol. (Lausanne). 4, 98.

Vinayagam, A., Kulkarni, M.M., Sopko, R., Sun, X., Hu, Y., Nand, A., Villalta, C., Moghimi, A., Yang, X., Mohr, S.E., et al. (2016). An Integrative Analysis of the InR/PI3K/Akt Network Identifies the Dynamic Response to Insulin Signaling. Cell Rep. 16, 3062–3074.

Ward, J.H. (1963). Hierarchical Grouping to Optimize an Objective Function. J. Am. Stat. Assoc. 58, 236–244.

Whiteman, E.L., Cho, H., and Birnbaum, M.J. (2002). Role of Akt/protein kinase B in metabolism. Trends Endocrinol. Metab. 13, 444–451.

Wolf, A., Rietscher, K., Glaβ, M., Hüttelmaier, S., Schutkowski, M., Ihling, C., Sinz, A., Wingenfeld, A., Mun, A., and Hatzfeld, M. (2013). Insulin signaling via Akt2 switches plakophilin 1 function from stabilizing cell adhesion to promoting cell proliferation. J. Cell Sci. 126, 1832–1844.

Workman, C.T., Yin, Y., Corcoran, D.L., Ideker, T., Stormo, G.D., and Benos, P. V. (2005). enoLOGOS: a versatile web tool for energy normalized sequence logos. Nucleic Acids Res. 33, W389–92.

Yugi, K., and Kuroda, S. (2017). Metabolism-Centric Trans-Omics. Cell Syst. 4, 19–20.

Yugi, K., Kubota, H., Toyoshima, Y., Noguchi, R., Kawata, K., Komori, Y., Uda, S., Kunida, K., Tomizawa, Y., Funato, Y., et al. (2014). Reconstruction of Insulin Signal Flow from Phosphoproteome and Metabolome Data. Cell Rep. 8, 1171–1183.

Yugi, K., Kubota, H., Hatano, A., and Kuroda, S. (2016). Trans-Omics: How To Reconstruct Biochemical Networks Across Multiple ?Omic? Layers. Trends Biotechnol. 34, 276–290.

Yusufi, F.N.K., Lakshmanan, M., Ho, Y.S., Loo, B.L.W., Ariyaratne, P., Yang, Y., Ng, S.K., Tan, T.R.M., Yeo, H.C., Lim, H.L., et al. (2017). Mammalian Systems Biotechnology Reveals Global Cellular Adaptations in a Recombinant CHO Cell Line. Cell Syst. 4, 530–542.e6.

Zhang, W., Thompson, B.J., Hietakangas, V., and Cohen, S.M. (2011). MAPK/ERK Signaling Regulates Insulin Sensitivity to Control Glucose Metabolism in Drosophila. PLoS Genet. 7, e1002429.

Zhang, Y., Zhang, Y., and Yu, Y. (2017). Global Phosphoproteomic Analysis of Insulin/Akt/mTORC1/S6K Signaling in Rat Hepatocytes. J. Proteome Res. acs.jproteome.7b00140.

Zimmet, P., Alberti, K.G.M.M., and Shaw, J. (2001). Global and societal implications of the diabetes epidemic. Nature 414, 782–787.

